# Coevolutionary Analysis and Perturbation-Based Network Modeling of the SARS-CoV-2 Spike Protein Complexes with Antibodies: Binding-Induced Control of Dynamics, Allosteric Interactions and Signaling

**DOI:** 10.1101/2021.01.19.427320

**Authors:** Gennady M. Verkhivker, Luisa Di Paola

**Affiliations:** Keck Center for Science and Engineering, Schmid College of Science and Technology, Chapman University, One University Drive, Orange, CA 92866, USA; Depatment of Biomedical and Pharmaceutical Sciences, Chapman University School of Pharmacy, Irvine, CA 92618, USA; Unit of Chemical-Physics Fundamentals in Chemical Engineering, Department of Engineering, Università Campus Bio-Medico di Roma, via Álvaro del Portillo 21, 00128 Rome, Italy

## Abstract

The structural and biochemical studies of the SARS-CoV-2 spike glycoproteins and complexes with highly potent antibodies have revealed multiple conformation-dependent epitopes highlighting the link between conformational plasticity of spike proteins and capacity for eliciting specific binding and broad neutralization responses. In this study, we used coevolutionary analysis, molecular simulations, and perturbation-based hierarchical network modeling of the SARS-CoV-2 S complexes with H014, S309, S2M11 and S2E12 antibodies targeting distinct epitopes to explore molecular mechanisms underlying binding-induced modulation of dynamics, stability and allosteric signaling in the spike protein trimers. The results of this study revealed key regulatory centers that can govern allosteric interactions and communications in the SARS-CoV-2 spike proteins. Through coevolutionary analysis of the SARS-CoV-2 spike proteins, we identified highly coevolving hotspots and functional clusters forming coevolutionary networks. The results revealed significant coevolutionary couplings between functional regions separated by the medium-range distances which may help to facilitate a functional cross-talk between distant allosteric regions in the SARS-CoV-2 spike complexes with antibodies. We also discovered a potential mechanism by which antibody-specific targeting of coevolutionary centers can allow for efficient modulation of allosteric interactions and signal propagation between remote functional regions. Using a hierarchical network modeling and perturbation-response scanning analysis, we demonstrated that binding of antibodies could leverage direct contacts with coevolutionary hotspots to allosterically restore and enhance couplings between spatially separated functional regions, thereby protecting the spike apparatus from membrane fusion. The results of this study also suggested that antibody binding can induce a switch from a moderately cooperative population-shift mechanism, governing structural changes of the ligand-free SARS-CoV-2 spike protein, to antibody-induced highly cooperative mechanism that can better withstand mutations in the functional regions without significant deleterious consequences for protein function. This study provides a novel insight into allosteric regulatory mechanisms of SARS-CoV-2 S proteins, showing that antibodies can modulate allosteric interactions and signaling of spike proteins, providing a plausible strategy for therapeutic intervention by targeting specific hotspots of allosteric interactions in the SARS-CoV-2 proteins.

## Introduction

The coronavirus disease 2019 (COVID-19) pandemic associated with the severe acute respiratory syndrome (SARS)^1–5^ has been at the focal point of biomedical research. SARS-CoV-2 infection is transmitted when the viral spike (S) glycoprotein binds to the host cell receptor leading to the entry of S protein into host cells and membrane fusion.^6–8^ The full-length SARS-CoV-2 S protein consists of two main domains, amino (N)-terminal S1 subunit and carboxyl (C)-terminal S2 subunit. The subunit S1 is involved in the interactions with the host receptor and includes an N-terminal domain (NTD), the receptor-binding domain (RBD), and two structurally conserved subdomains (SD1 and SD2). Structural and biochemical studies have shown that the mechanism of virus infection may involve spontaneous conformational transformations of the SARS-CoV-2 S protein between a spectrum of closed and receptor-accessible open forms, where RBD continuously switches between “down” and “up” positions where the latter can promote binding with the host receptor ACE2.^9–11^ The crystal structures of the S-RBD in the complexes with human ACE2 enzyme revealed structurally conserved binding mode shared by the SARS-CoV and SARS-CoV-2 proteins in which an extensive interaction network is formed by the receptor binding motif (RBM) of the RBD region.^12–16^ These studies established that binding of the SARS-CoV-RBD to the ACE2 receptor can be a critical initial step for virus entry into target cells. The rapidly growing body of cryo-EM structures of the SARS-CoV-2 S proteins detailed distinct conformational arrangements of S protein trimers in the prefusion form that are manifested by a dynamic equilibrium between the closed (“RBD-down”) and the receptor-accessible open (“RBD-up”) form required for the S protein fusion to the viral membrane.^17–26^ The cryo-EM characterization of the SARS-CoV-2 S trimer demonstrated that S protein may populate a spectrum of closed states by fluctuating between structurally rigid locked-closed form and more dynamic, closed states preceding a transition to the fully open S conformation.^26^ Structural and biophysical studies employed protein engineering to generate prefusion-stabilized SARS-CoV-2 S variants by introducing disulfide bonds and proline mutations to modulate stability of the S2 subunit and the inter-subunit boundaries, and consequently prevent refolding changes that accompany acquisition of the postfusion state.^27^ By combining targeted mutagenesis and cryo-EM structure determination, recent biophysical investigations demonstrated that modifications in the contact regions between the RBD and S2 domains via S383C/D985C double mutation can lead to the thermodynamic prevalence of the closed-down conformation, while the quadruple mutant (A570L/T572I/F855Y/N856I) perturbing the inter-protomer contacts can shift the equilibrium towards the open form with the enhanced binding propensities for the ACE2 host receptor.^28^ Protein engineering and cryo-EM studies of a prefusion-stabilized SARS-CoV-2 S ectodomain trimer using the inter-protomer disulfide bonds (S383C/D985C, G413C/P987C, T385C/T415C) between RBD and S2 regions can lock the trimer in the closed state and enhance the SARS-CoV-2 S resistance to proteolysis.^29^ Targeted design of thermostable SARS-CoV-2 spike trimers further specified how disulfide-bonded S-protein trimer variants imposing stabilization in the strategically located inter-protomer positions (S383-D385) and (G413-V987) can promote dramatic thermodynamic shifts towards the prefusion closed states with only ∼20% of the population corresponding to the open state.^30^ Recent biochemical studies of the SARS-CoV-2 S mutants with the enhanced infectivity profile^39–41^ discovered that a highly active D614G mutation can exert its dramatic functional effect on virus infectivity by radically shifting the population of the SARS-CoV-2 S trimer towards open states.^31^ The cryo-EM and sophisticated tomography tools determined the high-resolution structure and distribution of S trimers in situ on the virion surface.^32^ These studies confirmed a general mechanism of population shifts between different functional states of the SARS-CoV-2 S trimers, suggesting that RBD epitopes can become stochastically exposed to the interactions with the host receptor ACE2. Biophysical analysis of SARS-CoV-2 S trimer on virus particles revealed four distinct conformational states for the S protein and a sequence of conformational transitions through an obligatory intermediate in which all three RBD domains in the closed conformations are oriented towards the viral particle membrane.^33^ Cryo-EM structural studies also mapped a mechanism of conformational events associated with ACE2 binding, showing that the compact closed form of the SARS-CoV-2 S protein becomes weakened after furin cleavage between the S1 and S2 domains, leading to the increased population of partially open states and followed by ACE2 recognition that can accelerate transformation to a fully open and ACE2-bound form priming the protein for fusion activation.^34^

The early biochemical studies of SARS S proteins with antibodies (Abs) suggested that RBD regions of S proteins contain multiple conformation-dependent epitopes capable of inducing potent neutralizing Ab responses, thus revealing the link between conformational heterogeneity of S proteins and capacity for eliciting binding with highly potent neutralizing Abs.^35^ Subsequently, it was shown that major neutralizing epitopes of SARS-CoV may have been preserved during cross-species transmission, and that RBD-targeted Abs have a potential for broad protection against both human and animal SARS-CoV variants.^36^ The SARS-CoV-2 S protein–targeting monoclonal antibodies (mAbs) with potent neutralizing activity are of paramount importance and are actively pursued as therapeutic interventions for COVID-19 virus.^37–40^ The rapidly growing structural studies of SARS-CoV and SARS-CoV-2–neutralizing Abs targeting the RBD have suggested potential mechanisms underlying inhibition of the association between the S protein and ACE2 host receptor. The early structure of SARS-CoV-RBD complex with a neutralizing Ab 80R showed that the epitope on the S1 RBD overlapped closely with the ACE2-binding site, suggesting that a direct interference mechanism may be responsible for the neutralizing activity.^41^ However, several SARS-CoV–specific neutralizing Abs such as m396, 80R, and F26G19 that block the RBM motif in the open S conformation did not exhibit a strong neutralizing activity against SARS-CoV-2 protein. The crystal structure of a neutralizing Ab CR3022 in the complex with the SARS-CoV-2 S-RBD revealed binding to a highly conserved epitope that is located away from the ACE2-binding site but could only be accessed when two RBDs adopt the “up” conformation.^42^ Subsequent structural and surface plasmon resonance studies confirmed that CR3022 binds the RBD of SARS-CoV-2 displaying strong neutralization by allosterically perturbing the interactions between the RBD regions and ACE2 receptor.^43^ The proposed neutralization mechanism of SARS-CoV-2 through destabilization of the prefusion S conformation can provide a resistance mechanism to virus escape which can be contrasted with Abs directly competing for the ACE2-binding site and often susceptible to immune evasion. Potent neutralizing Abs from COVID-19 patients examined through electron microscopy studies confirmed that the SARS-CoV-2 S protein features multiple distinct antigenic sites, including RBD-based and non-RBD epitopes.^44^ These studies also suggested that some Abs may function by allosterically interfering with the host receptor binding and causing conformational changes in the S protein that can obstruct other epitopes and block virus infection without directly interfering with ACE2 recognition. Cryo–EM characterization of the SARS-CoV-2 S trimer in complex with the H014 Fab fragment revealed a new conformational epitope that is accessible only when the RBD is in the up conformation.^45^ Biochemical and virological studies demonstrated that H014 prevents attachment of SARS-CoV-2 to the host cell receptors and can exhibit broad cross-neutralization activities by leveraging conserved nature of the RBD epitope and a partial overlap with ACE2-binding region. The recently reported mAb S309 potently neutralizes both SARS-CoV-2 and SARS-CoV through binding to a conserved RBD epitope which is distinct from the RBM region and accessible in both open and closed states, so that there is no completion between S309 and ACE2 for binding to the SARS-CoV-2 S protein.^46^ Two ultra-potent Abs S2M11 and S2E12 targeting the overlapping RBD epitopes were recently reported, revealing Ab-specific modulation of protein responses and adaptation of different functional states for the S trimer.^47^ Cryo-EM structures showed that S2M11 can recognize and stabilize S protein in the closed conformation by binding to a quaternary epitope spanning two RBDs of the adjacent protomers in the S trimer, while S2E12 binds to a tertiary epitope contained within one S protomer and shifts the conformational equilibrium towards a fully open S trimer conformation.^47^ The mAbs isolated from 10 convalescent COVID-19 patients showed neutralizing activities against authentic SARS-CoV-2, with the mAb 4A8 displaying high potency by binding to the NTD of the S protein conformation with one RBD in “up” conformation and the other two RBDs in “down” conformation.^48^ Interestingly, none of the isolated mAbs recognize the RBD and inhibit binding of SARS-CoV-2 S protein to ACE2, suggesting that allosterically regulated mechanisms may underlie the functional effects and experimentally observed efficient cross-neutralization.^48^ Moreover, it was proposed that combining NTD-targeting 4A8 with RBD-targeting Abs may help in the design of “cocktail” therapeutics to combat the escaping mutations of the virus.

The most recent investigation reported discovery of an ultra-potent synthetic nanobody Nb6 that neutralizes SARS-CoV-2 by stabilizing the fully inactive down S conformation preventing binding with ACE2 receptor.^49^ Affinity maturation and structure-guided design produced a trivalent nanobody, mNb6-tri that simultaneously binds to all three RBDs, yielding the remarkably high affinity for S protein and completely blocking the S-ACE2 interactions by occupying the binding site and locking spike protein in the inactive, receptor-inaccessible state. In general, Abs tend to bind to the most easily accessible regions of the virus, where viruses can tolerate mutations and thereby escape immune challenge. The emerging body of recent studies suggested that properly designed cocktails of Abs can provide a broad and efficient cross-neutralization effects through synergistic targeting of conserved and more variable SARS-CoV-2 RBD epitopes, thereby offering a robust strategy to combat virus resistance.^45–49^

Computational modeling and molecular dynamics (MD) simulations have been instrumental in predicting conformational and energetic mechanisms of SARS-CoV-2 functions.^50–55^ Microsecond, all-atom MD simulations of the full-length SARS-CoV-2 S glycoprotein embedded in the viral membrane, with a complete glycosylation profile were recently reported, providing the unprecedented level of details about open and closed structures.^51^ MD simulations of the SARS-CoV-2 spike glycoprotein identified differences in flexibility of functional regions that may be important for modulating the equilibrium changes and binding to ACE2 host receptor.^52^ Computational studies examined SARS-CoV-2 S trimer interactions with ACE2 enzyme using the recent crystal structures^53–62^ providing insights into the key determinants of the binding affinity and selectivity. A comprehensive study employed MD simulations to reveal a balance of hydrophobic interactions and elaborate hydrogen-bonding network in the SARS-CoV-2-RBD interface.^59^ Molecular mechanisms of the SARS-CoV-2 binding with ACE2 enzyme were analyzed in our recent study using coevolution and conformational dynamics.^62^ Using protein contact networks and perturbation response scanning based on elastic network models, we recently discovered existence of allosteric sites on the SARS-CoV-2 spike protein.^63^ By using molecular simulations and network modeling we recently presented the first evidence that the SARS-CoV-2 spike protein can function as an allosteric regulatory engine that fluctuates between dynamically distinct functional states.^64^

In this study, we used a battery of computational approaches to explore and simulate molecular mechanisms underlying responses of the SARS-CoV-2 S proteins to binding of a panel of Abs (H014, S309, S2M11 and S2E12) that target distinct epitopes in the RBD regions. Using coevolutionary analysis, molecular simulations, and perturbation-based hierarchical network modeling of the SARS-CoV-2 S complexes with these Abs, we examined binding-induced modulation of dynamics, stability and allosteric interactions in the S protein trimers. The results of this study revealed structural topography of coevolutionary couplings and network connectivity that may determine mechanisms of allosteric signaling in the SARS-CoV-2 S proteins. We show that specific Ab targeting of conserved centers and coevolutionary hotspots in the S protein that are distinct from RBM region can allow not only for productive binding, but also for efficient Ab-induced modulation of long-range interactions between the S1 and S2 units of the SARS-CoV-2 S protein. Using perturbation-based network modeling, we find that targeted binding of the Abs could leverage direct contacts with coevolutionary hotspots to effectively restore allosteric potential of the S1 regions in the open states, thereby strengthening the allosteric interaction network and protecting the S protein machinery from dissociation of S1 subunit required for membrane fusion. The results of this study provide a novel insight into allosteric regulatory mechanisms of SARS-CoV-2 S proteins showing that the examined Abs can uniquely modulate signal communication providing a plausible strategy for therapeutic intervention by targeting specific hotspots of allosteric interactions in the SARS-CoV-2 proteins.

## Materials and Methods

### Sequence Conservation and Coevolutionary Analyses

Multiple sequence alignment (MSA) was obtained using the MAFFT approach^65^ and homologues were obtained from UNIREF90.^66, 67^ We employed Kullback-Leibler (KL) sequence conservation score *KLConsScore* using MSA profiles generated by hidden Markov models in Pfam database for the SARS-CoV S glycoproteins.^68, 69^ Three Pfam domains were utilized corresponding to S1, the NTD (*bCoV_S1_N*, Betacoronavirus-like spike glycoprotein S1, N-terminal, Pfam:PF16451, Uniprot SPIKE_CVHSA, pdb id 6CS0, residues 33-324), the RBD (*bCoV_S1_RBD,* Betacoronavirus spike glycoprotein S1, receptor binding, Pfam:PF09408, Uniprot SPIKE_CVHSA, pdb id 6CS0, residues 335-512) and the new C-terminal domain, CTD (*CoV_S1_C* Coronavirus spike glycoprotein S1, C-terminal. Pfam:PF19209, Uniprot SPIKE_CVHSA, pdb id 6CS0, residues 522-580). S2 is described in the family Pfam:PF01601 (Uniprot SPIKE_CVHSA, pdb id 6CS0, residues 622-1120) which contains an additional S2′ cleavage site, a fusion peptide, internal fusion peptide, heptad repeat 1/2 domains, and the transmembrane domain. The following Uniprot entries were used for sequence analysis : P59594: SPIKE_SARS (previously SPIKE_CVHSA) (pdb id 6CS0) and P0DTC2: SPIKE_SARS2 (pdb id 6VXX, 6VYB).

The KL conservation is calculated according to the following formula:

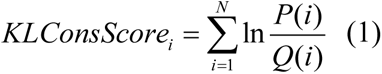

Here, *P*(*i*) is the frequency of amino acid *i* in that position and *Q*(*i*) is the background frequency of the amino acid in nature calculated using an amino acids background frequency distribution obtained from the UniProt database.^70^ To evaluate coevolutionary couplings in the SARS-CoV-2 S glycoproteins, we used MISTIC approach^71–73^ in which computation of residue covariation was done with three different direct coupling analysis (DCA) methods: mean field DCA (mfDCA),^74–76^ pseudo-likelihood maximization DCA (plmDCA)^77, 78^ and multivariate Gaussian modeling DCA (gaussianDCA).^79, 80^ For each residue, we computed cumulative covariation score (CScore) parameter, that evaluates to what degree a given position participates in the coevolutionary network. CScore is a derived score per position that characterizes the extent of coevolutionary couplings shared by a given residue. This score is calculated as the sum of covariation scores above a certain threshold (typically top 5% of the covariation scores) for every position pair where the particular position appears.

### Coarse-Grained Molecular Simulations

Coarse-grained (CG) models are computationally effective approaches for simulations of large systems over long timescales. In this study, CG-CABS model^81–85^ was used for simulations of the cryo-EM structures of the SARS-CoV-2 S complexes with H014, S309, S2M11, and S2E12 Abs. In this model, the amino acid residues are represented by Cα, Cβ, the center of mass of side chains and another pseudoatom placed in the center of the Cα-Cα pseudo-bond.^81–83^ We employed CABS-flex approach that efficiently combines a high-resolution coarse-grained model and efficient search protocol capable of accurately reproducing all-atom MD simulation trajectories and dynamic profiles of large biomolecules on a long time scale.^81–85^ The sampling scheme of the CABS model used in our study is based on Monte Carlo replica-exchange dynamics and involves a sequence of local moves of individual amino acids in the protein structure as well as moves of small fragments.^81–83^ CABS-flex standalone package dynamics implemented as a Python 2.7 object-oriented package was used for fast simulations of protein structures.^85^ A total of 1,000 independent CG-CABS simulations were performed for each of the studied systems. In each simulation, the total number of cycles was set to 10,000 and the number of cycles between trajectory frames was 100. The cryo-EM structures of the SARS-CoV-2 S trimer complexes with a panel of Abs including H014, S309, S2M11, and S2E12 were used in CG-CABS simulations (Figures 1,2). These structures included the partially open and fully open forms of the SARS-CoV-2 S trimer in the complex with H014 (Figure 1A,B), the partially closed and fully closed S trimer forms bound with S309 (Figure 1C,D), the fully closed S trimer form complexed with S2M11 (Figure 1E), and the fully open S trimer form in the complex with S2E12 (Figure 1F). All structures were obtained from the Protein Data Bank.^86, 87^ During structure preparation stage, protein residues in the crystal structures were inspected for missing residues and protons. Hydrogen atoms and missing residues were initially added and assigned according to the WHATIF program web interface.^88, 89^ The structures were further pre-processed through the Protein Preparation Wizard (Schrödinger, LLC, New York, NY) and included the check of bond order, assignment and adjustment of ionization states, formation of disulphide bonds, removal of crystallographic water molecules and co-factors, capping of the termini, assignment of partial charges, and addition of possible missing atoms and side chains that were not assigned in the initial processing with the WHATIF program. The missing loops in the cryo-EM structures were also reconstructed using template-based loop prediction approaches ModLoop^90^ and ArchPRED^91^ The conformational ensembles were also subjected to all-atom reconstruction using PULCHRA method^92^ and CG2AA tool^93^ to produce atomistic models of simulation trajectories. The protein structures were then optimized using atomic-level energy minimization with a composite physics and knowledge-based force fields as implemented in the 3Drefine method.^94^

**Figure 1.**
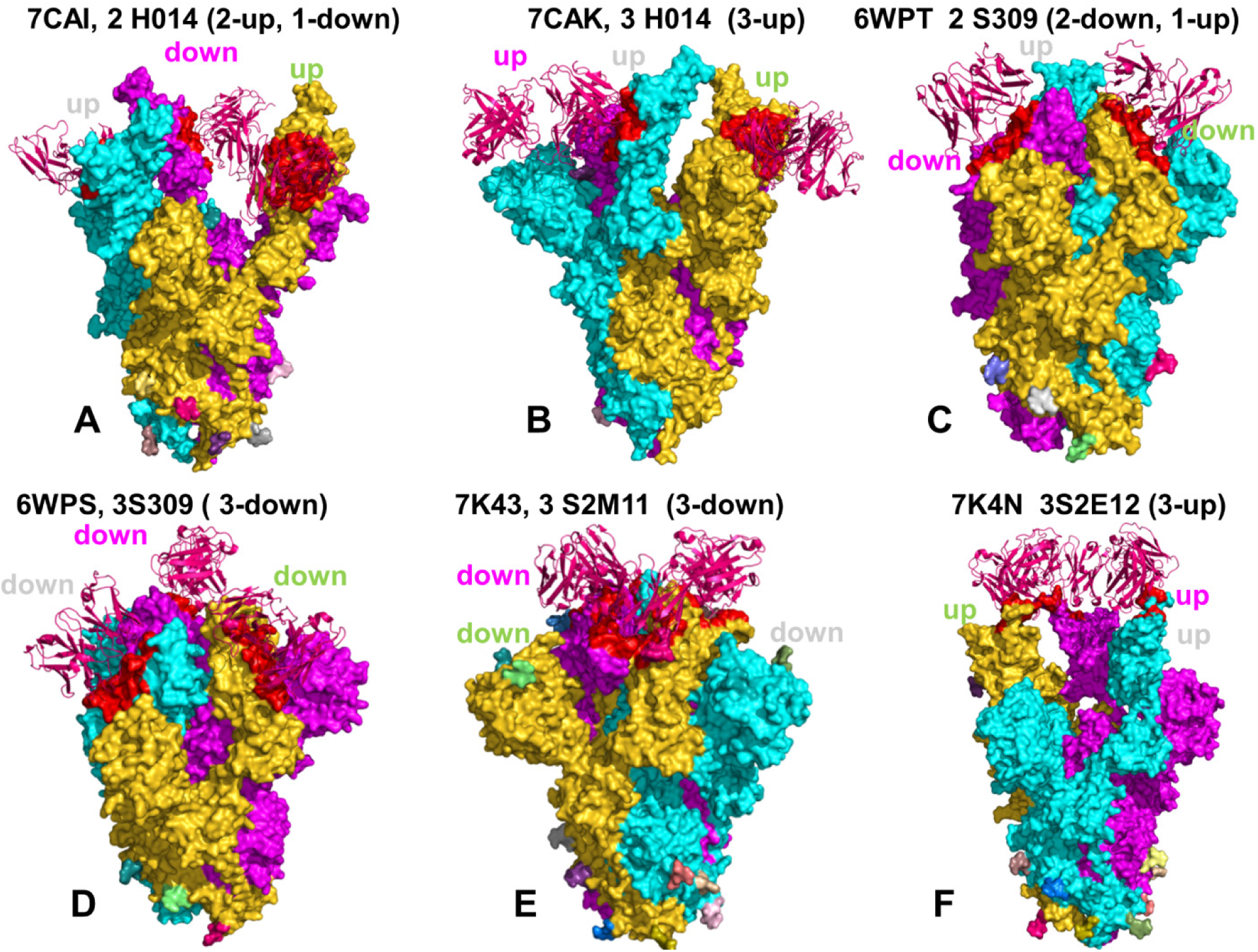
The cryo-EM structures of the SARS-CoV-2 S protein trimer complexes with a panel of Abs used in this study. (A) The cryo-EM structure of the SARS-CoV-2 S protein trimer with two RBDs in the open state complexed with two H014 Fab (pdb id 7CAI).^45^ (B) The cryo-EM structure of the SARS-CoV-2 S protein trimer with three RBD in the open state complexed with three H014 Fab (pdb id 7CAK).^45^ (C) The cryo-EM structure of the SARS-CoV-2 S protein trimer with two RBDs in the closed form and one RBD in the open state bound with the two S309 neutralizing Fab fragments (pdb id 6WPT).^46^ (D) The cryo-EM structure of the SARS-CoV-2 S protein trimer with all three RBDs in the closed form bound with the three S309 neutralizing Fab fragments (pdb id 6WPS).^46^ (E) The cryo-EM structure of the SARS-CoV-2 S protein trimer with all three RBDs in the closed-down form bound with the three S2M11 neutralizing Fab fragments (pdb id 7K43).^47^ (F) The cryo-EM structure of the SARS-CoV-2 S protein trimer with all three RBDs in the open-up form bound with the three S2ME12 neutralizing Fab fragments (pdb id 7K4N).^47^ The SARS-CoV-2 S proteins are shown in surface representation, with protomer A in green, protomer B in cyan, and protomer C in magenta. The Ab structures are shown in ribbons and colored in maroon. All structures are annotated and open/closed (up/down) conformations of S protomers are indicated.

**Figure 2.**
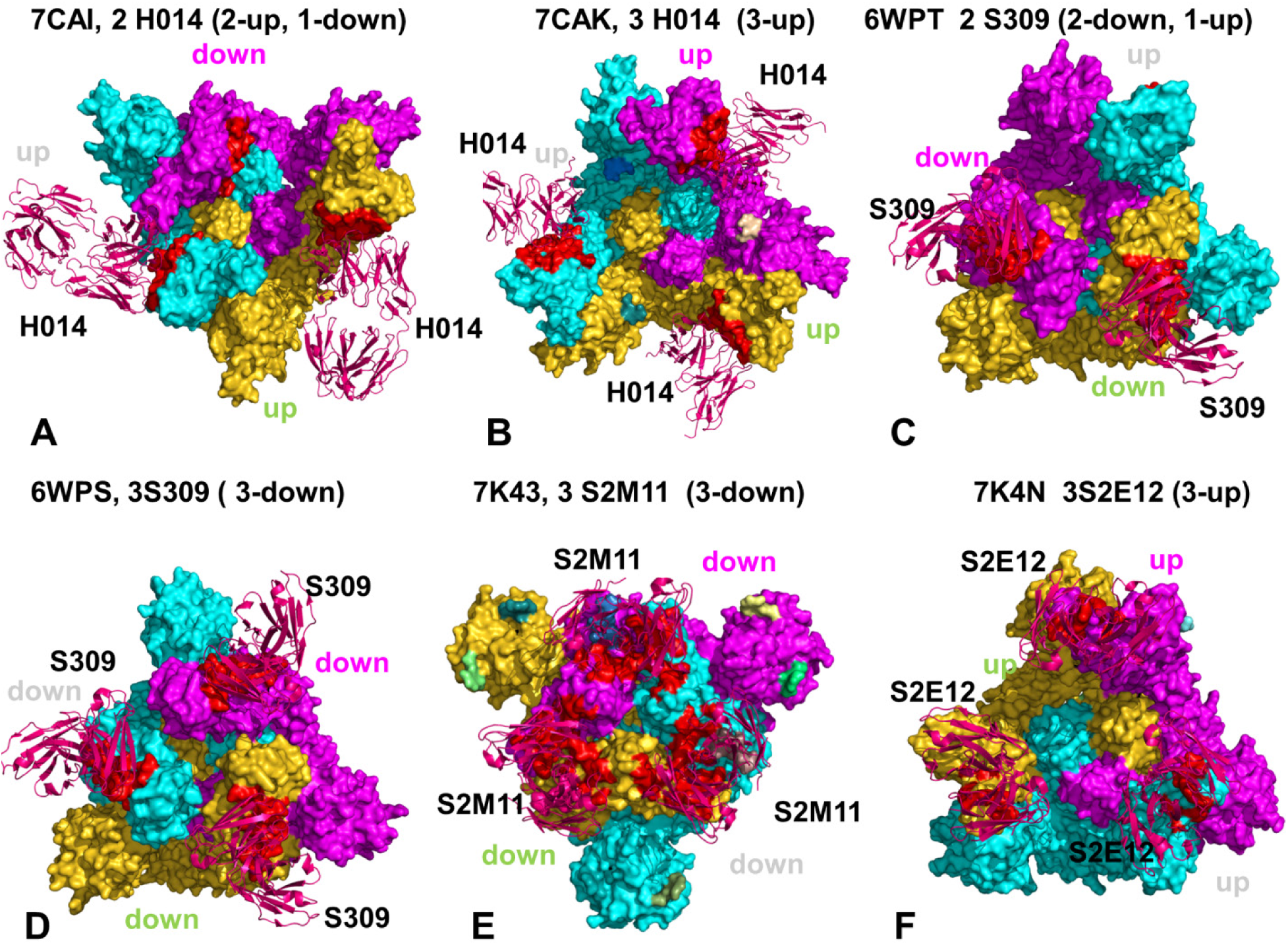
The binding epitopes of the SARS-CoV-2 S protein trimer complexes with a panel of Abs used in this study. The top view highlighting the binding epitopes is shown for the cryo-EM structure of the SARS-CoV-2 S protein trimer with H014 (A,B), S309 (C,D), S2M11 (E), an S2E12 (F). Note, S2M11 recognizes a quaternary epitope comprising distinct regions of two neighboring RBDs within an S trimer (E). The SARS-CoV-2 S proteins are shown in surface representation, with protomer A in green, protomer B in cyan, and protomer C in magenta. The Ab structures are shown in ribbons and colored in maroon. All structures are annotated and open/closed (up/down) conformations of S protomers are indicated.

### Protein Stability and Mutational Scanning Analysis

To compute protein stability changes in the SARS-CoV-2 trimer mutants, we conducted a systematic alanine scanning of protein residues in the SARS-CoV-2 trimer mutants. BeAtMuSiC approach was employed that is based on statistical potentials describing the pairwise inter-residue distances, backbone torsion angles and solvent accessibilities, and considers the effect of the mutation on the strength of the interactions at the interface and on the overall stability of the complex.^95–97^ The binding free energy of protein-protein complex can be expressed as the difference in the folding free energy of the complex and folding free energies of the two protein binding partners:

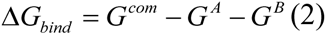

The change of the binding energy due to a mutation was calculated then as the following:

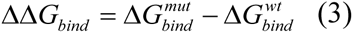

We leveraged rapid calculations based on statistical potentials to compute the ensemble-averaged binding free energy changes using equilibrium samples from MD trajectories. The binding free energy changes were computed by averaging the results over 1,000 equilibrium samples for each of the studied systems.

### Protein Contact Networks and Network Clustering

The protein contact network is a network whose nodes are the protein residues and links are active contacts between residues in the protein structure. The protein contact network is an undirected, unweighted graph; it is built on the basis of the distance matrix **d**, whose generic element *d_ij_* records the Euclidean distance between the *i*th and the *j*th residue (measured between the corresponding α carbons). A detailed description of network construction and significance of network descriptors is presented in our previous studies.^98–100^ The active network links are defined using a range of contacts between 4 Å and 8 Å. The description of the network is given by the following adjacency matrix :

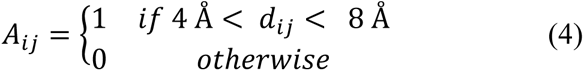

Where *d_i,j_* is the distance between the residues. The node degree describes the number of links each residue has with other residues, defined as:

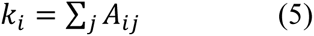

We previously demonstrated that spectral network clustering targets functional modules in proteins.^98^ The network clustering is based on the spectral decomposition of the network Laplacian

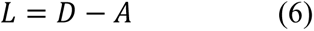

where *D* is the degree matrix, a diagonal matrix whose diagonal is the degree vector, and *A* is the adjacency matrix. We used the eigenvector corresponding to the second minor eigenvalue *v*_2_ of the Laplacian (Fiedler’s vector) to assign nodes (residues) to different clusters. We introduced a novel feature in the hierarchical binary algorithm to compute any number of clusters *n_clus_* (no more only powers of two): we parted the whole range of values of *v*_2_ into *n_clus_* parts, of length *l_clus_*; so residues are assigned to the first cluster if the corresponding component falls between min(***v***_2_) and min(***v***_2_) + *l_clus_* the generic i-th cluster, thus, is that made of residues corresponding to ***v***_2_ components comprised in the range [min(***v***_2_) + (*i* − 1) · *l_clus_*, min(***v***_2_) + *i* · *l_clus_*].

Once the network is divided into a given number of clusters (powers of two), we define the participation coefficient, defined as:

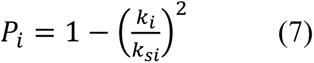

where *k_i_* is the overall node degree, while *k_si_* is the node degree including only links with nodes (residues) that belong to their own cluster. The participation coefficient *P* describes the propensity of residue nodes to participate into inter-cluster communication. We designate as highly active communication residues the nodes with *P>0.75*, based on our previous studies showing that such residues typically correspond to important regulatory centers of signal transmission between protein domains.^98^ The proposed methodology of network clustering was implemented as Cytoscape plugin.^101^

In the framework of hierarchical network modeling approach, we also employed a graph-based representation of protein structures^102–104^ with residues as network nodes and the inter-residue edges as residue interactions to construct the residue interaction networks using dynamic correlations^104^ and coevolutionary residue couplings ^105^ as detailed in our previous studies.^105–107^ The ensemble of shortest paths is determined from matrix of communication distances by the Floyd-Warshall algorithm.^108^ Network graph calculations were performed using the python package NetworkX.^109^ Using the constructed protein structure networks, we computed the residue-based betweenness parameter. The short path betweenness centrality of residue *i* is defined to be the sum of the fraction of shortest paths between all pairs of residues that pass through residue *i* :

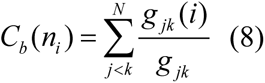

where *g_jk_* denotes the number of shortest geodesics paths connecting *j* and *k,* and *g_jk_*(*i*) is the number of shortest paths between residues *j* and *k* passing through the node *n_i_* .

### Perturbation Response Scanning

Perturbation Response Scanning (PRS) approach^110, 111^ was used to estimate residue response to external forces applied systematically to each residue in the protein system. This approach has successfully identified hotspot residues driving allosteric mechanisms in single protein domains and large multi-domain assemblies.^112–117^ The implementation of this approach follows the protocol originally proposed by Bahar and colleagues^112, 113^ and was described in details in our previous studies.^64^ In brief, through monitoring the response to forces on the protein residues, the PRS approach can quantify allosteric couplings and determine the protein response in functional movements. In this approach, it 3N × 3*N* Hessian matrix ***H*** whose elements represent second derivatives of the potential at the local minimum connect the perturbation forces to the residue displacements. The 3*N*-dimensional vector ***ΔR*** of node displacements in response to 3*N*-dimensional perturbation force follows Hooke’s law ***F*** = ***H*** * ***ΔR***. A perturbation force is applied to one residue at a time, and the response of the protein system is measured by the displacement vector ***ΔR***(*i*) = ***H***^−1^***F***^(i)^ that is then translated into *N*×*N* PRS matrix. The second derivatives matrix ***H*** is obtained from simulation trajectories for each protein structure, with residues represented by *C*_α_ atoms and the deviation of each residue from an average structure was calculated by Δ**R***_j_* (*t*) = **R**_j_ (*t*) − 〈**R**_j_(*t*)〉, and corresponding covariance matrix C was then calculated by Δ**R**Δ**R***^T^*. We sequentially perturbed each residue in the SARS-CoV-2 spike structures by applying a total of 250 random forces to each residue to mimic a sphere of randomly selected directions.^64^ The displacement changes, Δ***R^i^*** is a *3N-*dimensional vector describing the linear response of the protein and deformation of all the residues. Using the residue displacements upon multiple external force perturbations, we compute the magnitude of the response of residue *k* as 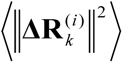 averaged over multiple perturbation forces **F**^(^*^i^*^)^, yielding the *ik*^th^ element of the *N*×*N* PRS matrix.^112, 113^ The average effect of the perturbed effector site *i* on all other residues is computed by averaging over all sensors (receivers) residues *j* and can be expressed as 〈(Δ***R^i^***)^2^〉_effector_. The effector profile determines the global influence of a given residue node on the perturbations in other protein residues and can be used as proxy for detecting allosteric regulatory hotspots in the interaction networks. In turn, the *j* ^th^ column of the PRS matrix describes the sensitivity profile of sensor residue *j* in response to perturbations of all residues and its average is denoted as 〈(Δ***R^i^***)^2^〉_sensor_. The sensor profile measures the ability of residue *j* to serve as a receiver (or transmitter) of dynamic changes in the system.

## Results and Discussion

### Sequence Analysis Links Evolutionary Patterns in SARS-CoV S Proteins with Antibody Binding Preferences

To determine the evolutionary patterns in the SARS-CoV S proteins and characterize the extent of conservation and variability of the S1 and S2 subunits, we utilized KL sequence conservation score.^71–73^ Consistent with previous studies^118–120^ we found that S1 RBD is less conserved than domains in the S2 subunit (Figure 3A,B). The S2 subunit contains an N-terminal hydrophobic fusion peptide (FP), fusion peptide proximal region (FPPR), heptad repeat 1 (HR1), central helix region (CH), connector domain (CD), heptad repeat 2 (HR2), transmembrane domain (TD) and cytoplasmic tail (CT). The S1 domains are situated above the S2 subunit, covering and protecting the fusion apparatus. The results confirmed the higher conservation of the S2 subunit particularly highlighting conservation of the HR1 (residues 910-985), CH (residues 986-1035), CD (residues 1068-1163), HR2 (residues 1163-1211), and TD regions (residues 1211-1234) (Figure 3). The S2 subunit regions are highly conserved in the SARS-CoV and the SARS-CoV-2 variants while the S1 subunit was more diverse in the NTD and RBD regions.

**Figure 3.**
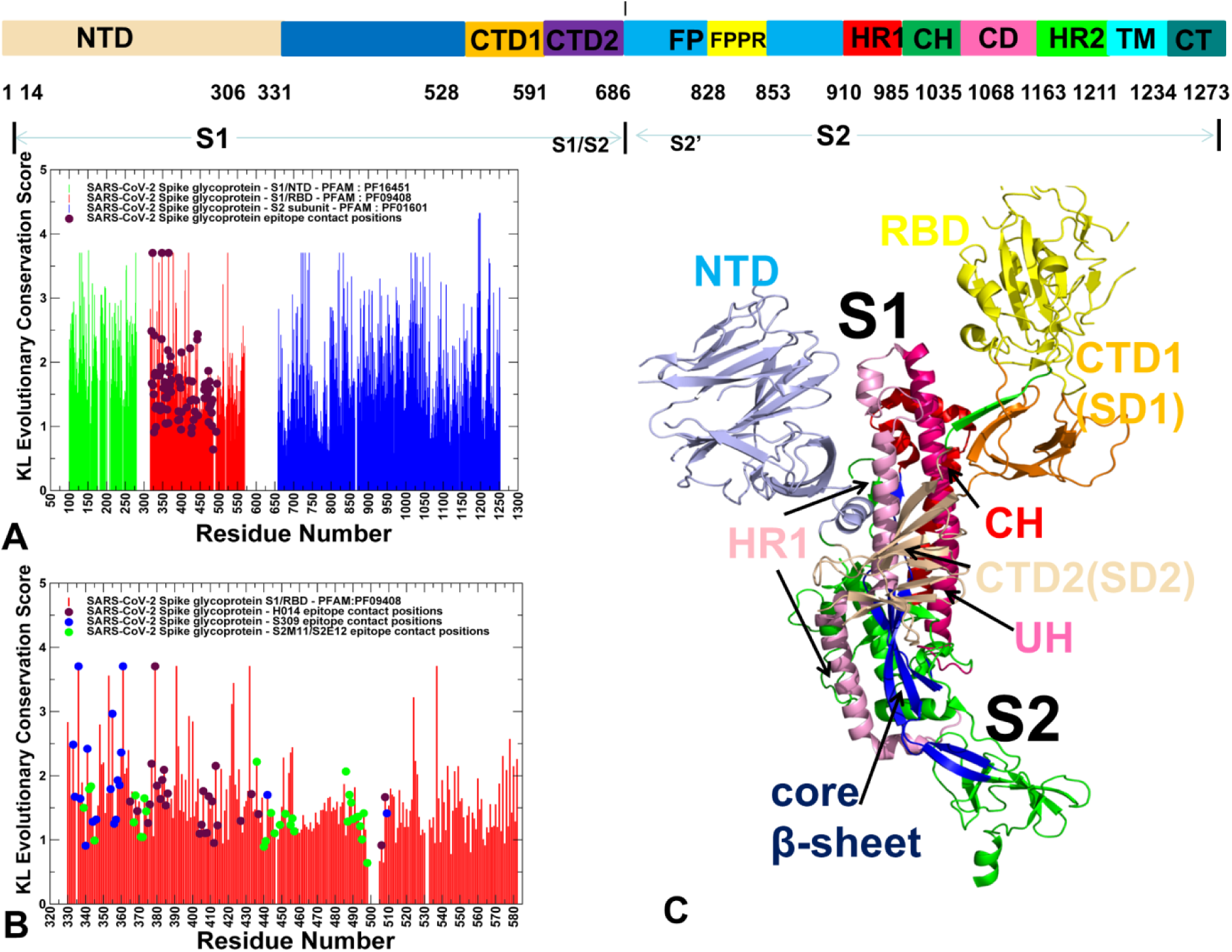
Sequence conservation analysis of the SARS-CoV-2 S glycoprotein. (Top panel) A schematic representation of domain organization and residue range for the full-length SARS-CoV-2 spike (S) protein. The subunits S1 and S2 include NTD RBD, C-terminal domain 1(CTD1), C-terminal domain 2 (CTD2), S1/S2 cleavage site (S1/S2), S2’ cleavage site (S2’), fusion peptide (FP), fusion peptide proximal region (FPPR), heptad repeat 1 (HR1), central helix region (CH), connector domain (CD), heptad repeat 2 (HR2), transmembrane domain (TM), and cytoplasmic tail (CT). (Panel A) The KL conservation score for SARS-CoV-RBD S protein. High KL scores indicate highly conserved sites and low scores correspond to more variable positions. Three Pfam domains were utilized corresponding to S1, the NTD (*bCoV_S1_N*, Betacoronavirus-like spike glycoprotein S1, N-terminal, Pfam:PF16451, Uniprot SPIKE_CVHSA, pdb id 6CS0, residues 33-324), the RBD (*bCoV_S1_RBD,* Betacoronavirus spike glycoprotein S1, receptor binding, Pfam:PF09408, Uniprot SPIKE_CVHSA, pdb id 6CS0, residues 335-512) and the new C-terminal domain, CTD (*CoV_S1_C* Coronavirus spike glycoprotein S1, C-terminal. Pfam:PF19209, Uniprot SPIKE_CVHSA, pdb id 6CS0, residues 522-580). S2 is described in the family Pfam:PF01601 (Uniprot SPIKE_CVHSA, pdb id 6CS0, residues 622-1120). The KL scores for the S1-NTD residues are shown in green bars, for the S1-RBD regions in the red bars, and for S2 residues in blue bars. The KL conservation scores for the epitope residues of all studied Abs are shown in filled maroon-colored circles. (Panel B) A close-up view of KL conservation scores for RBD regions of the SARS-CoV-2 S protein (Pfam:PF09408, Uniprot P0DTC2: SPIKE_SARS2 (pdb id 6VXX, 6VYB residue numbering) is shown in red bars. The KL scores are highlighted for the binding epitope residues of H014 (filled maroon-colored circles), S309 (filled blue circles), and S2M11/S2E12 (filled green circles). (Panel C) The structural organization of the SARS-CoV-2 S protein major domains is shown for a single protomer. The subunits S1 regions are annotated as follows : NTD (residues 14-306) in light blue; RBD (residues 331-528) in yellow; CTD1 (residues 528-591) in orange; CTD2 (residues 592-686) in wheat color ; upstream helix (UH) (residues 736-781) in red; HR1 (residues 910-985) in pink; CH (residues 986-1035) in hot pink; antiparallel core β-sheet (residues 711-736, 1045-1076) (in blue).

Among most conserved residues in the S2 subunit are clusters of conserved cysteine residues forming disulfide bridges that are crucial for stabilization of both pre-fusion and post-fusion SARS-CoV-2 spike protein conformation. The S proteins can contain up to 40 cysteine residues, 36 of which are conserved in the S proteins of various SARS-coronaviruses.^121^ The conserved cysteine cluster in the TD region 1220-CCMTSCCSC-1228 displayed high conservation scores, with C1121, M1222, and C1225 featuring top 1% of conservation scores (Figure 3A). Indeed, mutagenesis of the cysteine cluster I (1220-CCMTS-24), located immediately proximal to the TD showed the 55% reduction in S-mediated cell fusion as compared to the wild-type S protein.^122^

The proximal cysteine cluster 1225-CCSC-1228 is similarly important as alanine mutations in this cluster resulted in the 60% reduction of S-mediated cell fusion.^122^ At the same time, the nearest cysteine-rich cluster 1230-CSCGSCCK-1237 featured only one highly conserved C1235. According to the experimental data, mutations in this region caused only a moderate 15% reduction in cell fusion^122^, indicating that functional role of these clusters may be closely linked with the conservation level of cysteine residues.

As cysteine residues play critical roles in structural stabilization of proteins, we focused our attention on these residues and their locations in the sequences. Interestingly, the most conserved S2 positions included cysteine residues C720, C725, C731, C742, C822, C822, C833, C1014,C1025, and C1064 (Figure 3A). A conserved region flanked by C822 and C833 is known to be important for interactions with components of the SARS-CoV S trimer to control the activation of membrane fusion.^123^ In addition, other conserved residues included Y819, I800, L803, D830, L831, Y855, H1030, P1039, H1046 (Figure 3A,C). This is consistent with the experimental mutagenesis study based on cell-cell fusion and pseudovirion infectivity assay showing a critical role of the core-conserved residues C822, D830, L831, and C833 residues.^123^ Some of these residues are located C-terminal to the SARS-CoV S2 cleavage site at R797 forming a highly conserved region 798-SFIEDLLFNKVTLADAGF-815 that plays an important role for membrane fusion.^124^ Among highly conserved S protein regions are also six clusters of cysteine residues in the S2 subunit forming disulfide bridges crucial for stabilization of both pre-fusion and post-fusion SARS-CoV-2 spike protein conformations^125–127^ (Figures 3,4).

**Figure 4.**
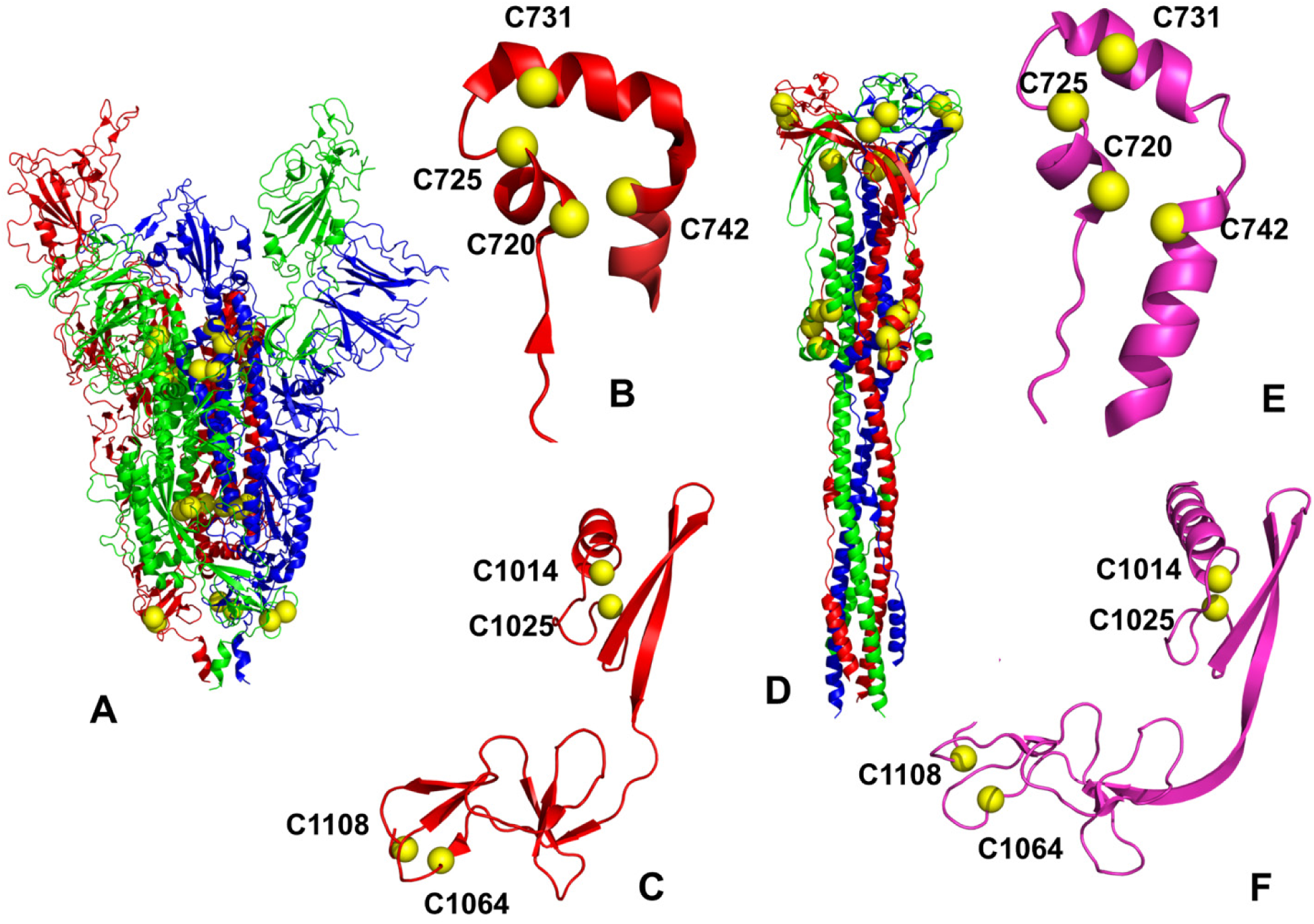
Sequence and structural conservation of cysteine clusters in the SARS-CoV-2 spike prefusion and postfusion states. (A) The cryo-EM structure of the SARS-CoV-2 S protein in the prefusion form is shown in ribbons. The protomer A is in green, protomer B in red, and protomer C in blue colors. The positions of the conserved cysteine clusters are shown in yellow spheres. (B) Structural arrangement of the conserved cluster formed by C720, C725, C731, and C742 in the UH region. The UH fragment is shown in red ribbons and conserved cysteine sites are shown in yellow spheres and annotated. (C) Structural organization of the conserved cysteine cluster in the β-hairpin region formed by C1014 and C1025, C1064 and C1108. The protein fragment is shown in red ribbons and conserved cysteines are in yellow spheres and annotated. (D) The cryo-EM structure of the SARS-CoV-2 S protein in the postfusion form. The protomer A is in green, protomer B in red, and protomer C in blue colors. The conserved cysteine clusters are in yellow spheres. (E) Structural arrangement of conserved cluster formed by C720, C725, C731, and C742 in the postfusion state. (F) Structural organization of a conserved cysteine cluster (C1014 and C1025, C1064 and C1108) in the post-fusion state. The conserved cysteine sites are shown in yellow spheres and annotated.

Structural analysis demonstrated that the core elements of S2 regions anchored by several cysteine clusters are highly preserved in the SARS-CoV-2 spike prefusion and postfusion states (Figure 4A,D) despite massive conformational changes of the SARS-CoV-2 S2 machinery.^125, 126^ Some of these regions include cysteine clusters formed by C720, C725, C731, and C742 in the upstream helix (UH) regions. (Figure 4B,E). Another conserved cysteine cluster of disulfide bonds is formed in the β-hairpin region (residues 1045-1076) located downstream of the CH region by residues C1014 and C1025 (C1032 and C1043 respectively in SPIKE_SARS2 sequence numbering) as well as residues C1064 and C1108 (C1082 and C1126 in SPIKE_SARS2 sequence numbering) (Figure 4C,F). This conserved segment of S2 subunit is a part of the antiparallel core β-sheet assembled from an N-terminal β-strand (β_46_) and a C-terminal β-hairpin (β_49_–β_50_). The top conserved RBD positions included C336, R355, C361, F374, F377, C379, L387, C391, D398, G413, N422, Y423, L425, F429, C432, and W436 (Figure 3B). The RBD region includes eight conserved cysteine residues, six of which form three disulfide linkages (C336–C361, C379– C432 and C391–C525), which stabilize the β-sheet RBD structure in the SARS-CoV-2 S protein. The crystal structures of S-proteins highlighted that two of these disulfide bonds are potentially redox-active, facilitating the primal interaction between the receptor and the spike protein.^121^

We particularly focused on conservation patterns of the SARS-CoV-2 RBD residues forming binding epitopes for H014, S309, S2E12 and S2M11 Abs. The H014 epitope is fairly large and broadly distributed across RBD regions (residues 368 to 386, 405 to 408 and 411 to 413, 439, and 503) forming a cavity on one side of the RBD (Tables S1 and S2, Supporting Information). Although most of contacts are formed with moderately conserved residues, H014 makes favorable interactions with two most highly conserved F377 and C379 positions in the RBD region (Figure 3B). Additionally, H014 makes strong interaction contacts with several other conserved RBD positions including Y380, S383, P384, K386, G413, and W436 (Figure 3B). Of particular importance are H014 contacts with S383 and G413 residues that are located at the inter-protomer boundaries (S383-D385) and (G413-V987) and could function as regulatory switches of S protein equilibrium.^30^ It is worth mentioning that disulfide-bonded S-protein trimer variants S383C/D985C at the RBD to S2 boundaries can lead to a predominant population of the prefusion closed states.^30^ Notably, the conformational epitope for H014 is only accessible when the RBD in an open conformation. According to our analysis, H014 interactions with conserved positions F377, C379, S383 located away from the RBM region could be important for binding and modulation of the enhanced cross-neutralization activities.

S309 engages an epitope distinct from the RBM making contacts with two most conserved RBD positions C336 and C361 (Figure 3B, Tables S3 and S4, Supporting Information). These residues form one of the disulfide linkages Cys336–Cys361 that stabilize the β-sheet RBD structure. S309 forms particularly strong contacts with neighboring residues L335 and P337 that displayed more moderate conservation. Our analysis indicated that the KL evolutionary score of the S309 contact RBD positons is considerably higher than average, with several interacting residues such as T333, C336, V341, R355, C361 displaying particularly strong conservation (Figure 3B). We suggest that binding to these highly conserved and structurally rigid positions located away from the ACE2-binding interface may contribute to a broad neutralizing activity of S309. Indeed, viruses may have evolved to maintain the sensitive regions of their structure inaccessible to the immune system. As a result, Abs tend to bind to the most easily accessible regions of the virus, where viruses can tolerate mutations. By targeting evolutionary conserved RBD epitopes H014 and S309 Abs can potentially better combat virus resistance. Based on this analysis, we hypothesized that binding of H014 and S309 to conserved epitopes of S proteins that are distinct from ACE2-binding site may involve allosterically regulated mechanism in which Abs induce long-range alterations in the interaction networks and allosteric communications.

S2M11 recognizes a quaternary epitope through electrostatic interactions and shape complementarity, comprising distinct regions of two neighboring RBDs within an S trimer.^47^ This Ab is believed to induce inhibition of membrane fusion through conformational trapping of SARS-CoV-2 S trimer in the closed state.^47^ S2M11 forms contacts with highly conserved sites F374 and W436 on one RBD and makes interactions with F486 on the RBM motif of the other RBD (Table S5, Supporting Information, and Figure 3B). We also characterized the inter-molecular contacts formed by S2E12 (Table S6, Supporting Information) that targets the RBM motif and can block the attachment to the ACE receptor. According to our analysis, the contact positions targeted by these Abs are only moderately conserved as the epitope overlaps with a more variable RBD region (Figure 3A,B).

### Coevolutionary Analysis of the SARS-CoV-2 Proteins Reveals Regulatory Centers and Functional Role of the Epitope Regions in the Network of Evolutionarily Coupled Residues

Coevolutionary dependencies of protein residues can mediate protein recognition and are often spatially close to each other, forming clusters of interacting residues that are located near functionally important sites.^129–132^ The coevolving residues may be clustered in mobile regions and form interaction networks of evolutionarily coupled residues that facilitate protein conformational changes.^133^ Using MISTIC approach^71–73^ we determined coevolutionary dependencies between S protein residues using plmDCA (Figure 5) and gaussianDCA models (Figure S1, Supporting Information). The network of coevolutionary couplings in the SARS-CoV-2 S structures was then constructed in which the nodes represented protein residues and links corresponded to coevolutionary dependencies between these residues. To identify critical nodes of this coevolutionary network that may coordinate and transmit coevolutionary signals, we computed plmDCA-based cScore profiles (Figure 5A) that measure the global influence of a given position in a coevolutionary network. This score is calculated as the sum of covariation scores above a certain threshold (top 5% of the covariation scores) for every position pair where the particular position appears. Using this approach, we quantified coevolutionary relationships between residues, identified coevolutionary couplings for functionally important regions and also mapped high CScore positions onto the binding epitopes for studied SARS-CoV-2 complexes (Figure 5B).

**Figure 5.**
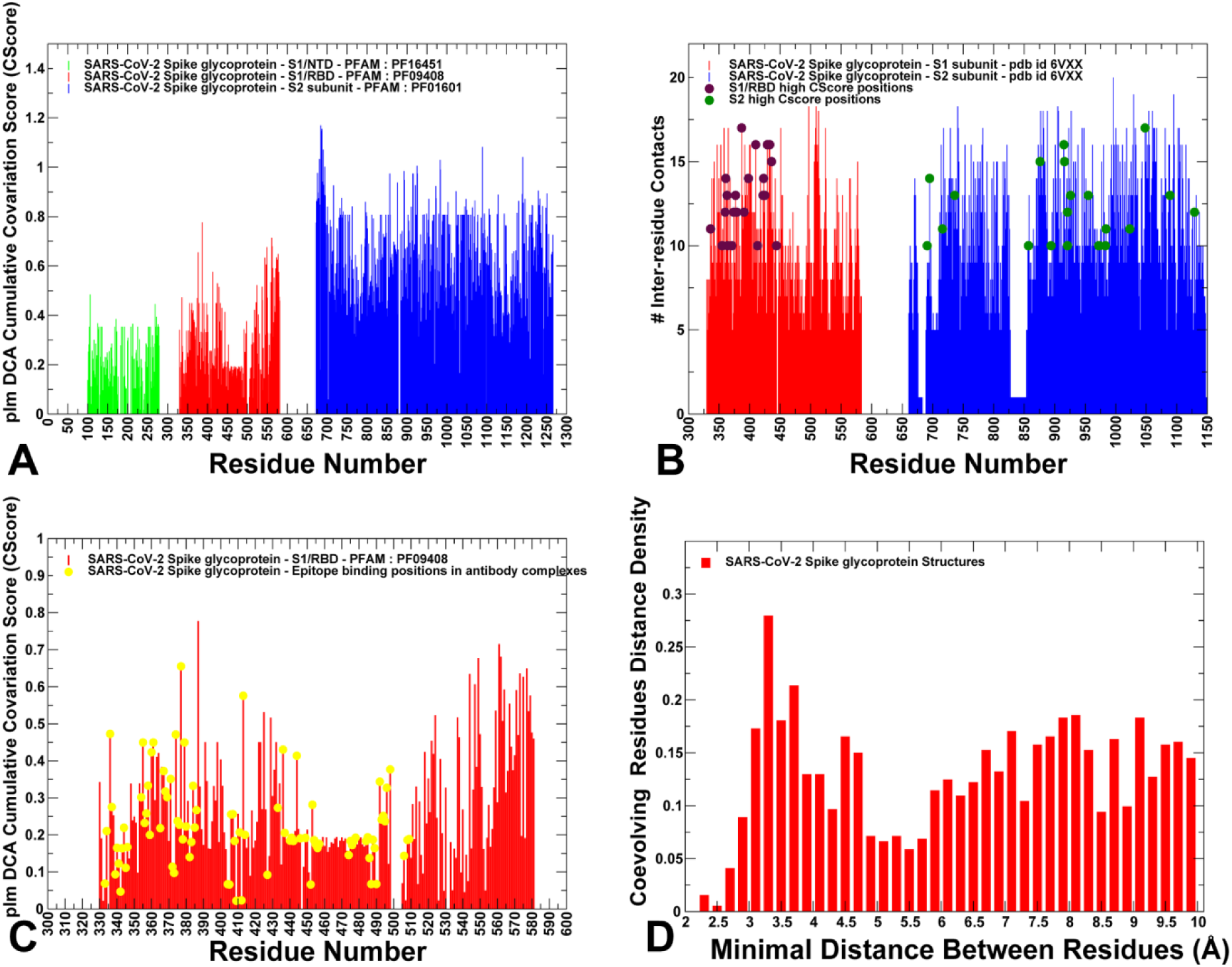
Coevolutionary profiles of the SARS CoV-2 S proteins. (A) The plmDCA-based coevolutionary Cscore profile for the SARS-CoV-2 S proteins (P0DTC2: SPIKE_SARS2 sequence numbering). The Cscore values are shown for the S1-NTD residues in green bars (Pfam:PF16451), for the RBD in red bars (Pfam:PF09408) and for S2 regions in blue bars (Pfam:PF01601). (B) A close-up of the CSscore profile for the RBD regions is shown in red bars. The CScores for the binding epitope residues of H014, S309, S2M11, and S2E12 are shown in filled green circles. (C) The distribution of the inter-residue contacts in the S1-RBD regions (red bars) and S2 regions (blue bars). The highly coevolving centers in the RBD regions are in maroon-colored filled circles and the high CSscore residues in S2 regions are in orange-colored filled circles. (D) The distance probability distribution of directly coupled residue pairs in the studied SARS-CoV-2 S complexes is shown in red filled bars.

We focused our analysis on the computed distribution of plmDCA-based CScore profiles (Figure 5A) that revealed several important trends. First, the results revealed an appreciable density of coevolving centers in the S1 subunit, primarily in the RBD and especially CTD1 regions. This pattern can be further illustrated by a circular representation of the pairwise coevolutionary scores (Figure S2, Supporting Information) showing the greater concentration of coevolutionary links anchored by the CTD1 regions (residues 528-591). The distribution of CScores pointed to the significantly higher density of coevolutionary couplings in the tightly packed S2 subunit (Figure 5A). The residues with significant CScore values are distributed across various S2 regions, including UH (residues 736-781), CH (residues 986-1035), HR1 region (residues 910-985), HR2 (residues 1163-1211) and β-hairpin (BH) region (residues 1035-1071). A dense network of coevolutionary coupled residues in the S2 regions can be therefore detected as evident from a graphical annotation of the pairwise coevolutionary scores (Figure S3, Supporting Information). Interestingly, the distribution of CScore values for the epitope residues showed that many contact positions are aligned with highly coevolving residues (Figure 5B). Among high cScore sites that establish intermolecular contacts with Abs are C336, R355, C361, F374, F377, C379, L387, C391, D398, G413, N422, Y423, L425, F429, C432, W436 (Figure 5B). Some of these residues are highly conserved (C336, C361, C379) while other sites exhibited moderate to high conservation level (L387, W436).

Structural analysis of coevolutionary hotspots corresponding to the local maxima of the distribution revealed presence of clusters situated in functional regions (Figure 6). We observed that coevolutionary centers can be localized in the key regions of the SARS-CoV-2 S protein, occupying the proximity of the SA1/S2 cleavage site, the HR1 and CH regions of S2 subunit as well as RBD and CTD1 regions in the S1 domain (Figure 6). Of particular interest are several coevolutionary hotspots located near a well-recognized cleavage site at the S1/S2 boundary. The furin cleavage site emerges as a disordered loop (residues 655-GICASYHTVSLLRST-669 in the SPIKE_SARS sequence or residues 669-GICASYQTQT-NSPRRARSVA-688 in SPIKE_SARS2 sequence).

**Figure 6.**
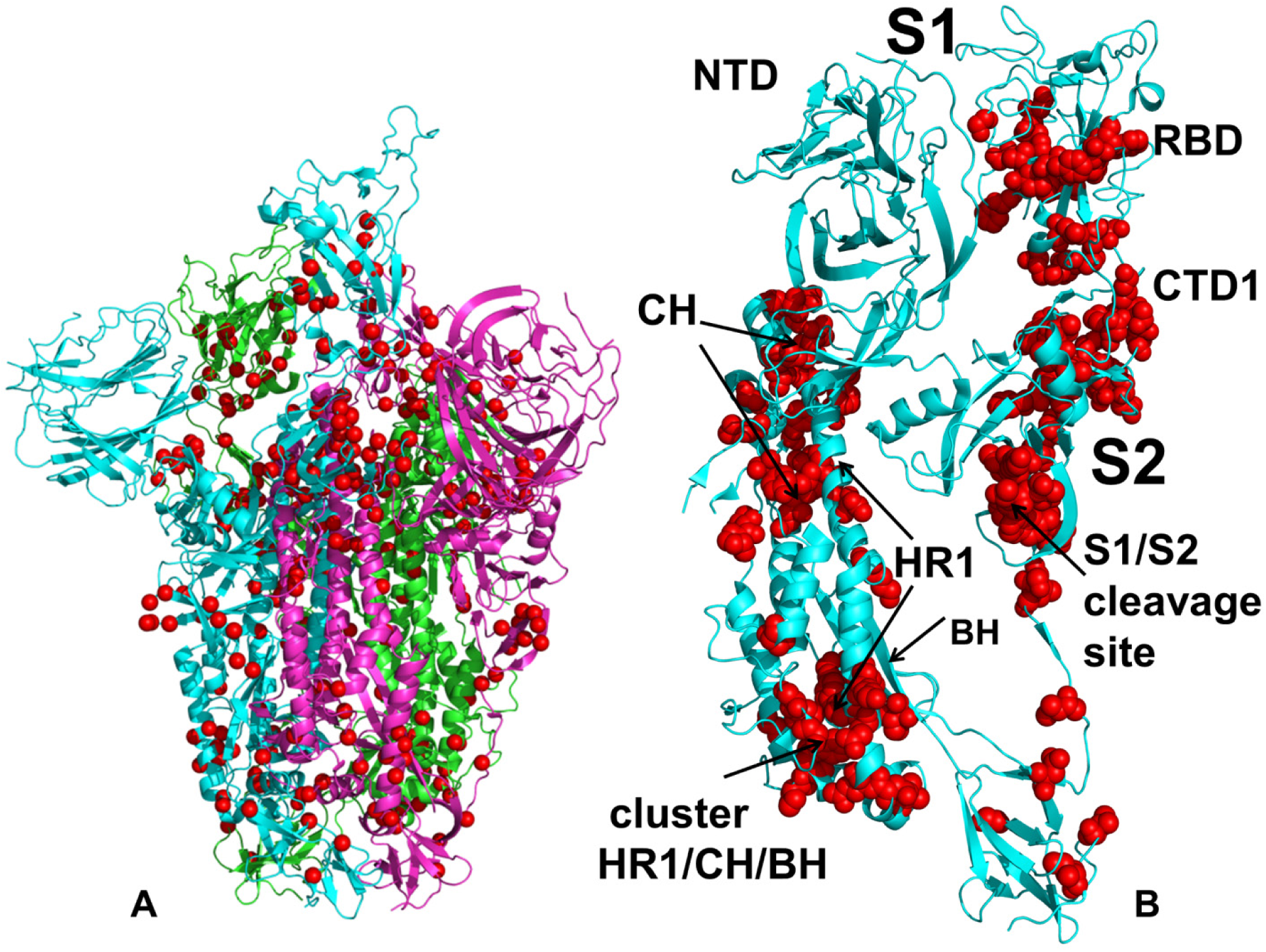
Structural analysis of coevolutionary hotspots in the SARS-CoV-2 S proteins. (A) Structural map of high CSscore residues shown in red spheres is projected on the cryo-EM structure of the SARS-CoV-2 S protein. (B) A close-up of coevolutionary centers mapped onto a single protomer of the S protein. The protomer is shown in cyan ribbons and high CSscore positions are depicted in red spheres. The map shows localization of coevolutionary hotspots in the key regions of the SARS-CoV-2 S protein, occupying the proximity of the SA1/S2 cleavage site, the HR1 and CH regions of S2 subunit as well as RBD and CTD1 regions in the S1 domain.

Our analysis showed that residues immediately C-terminal to the S1/S2 cleavage site such as S670, K672, S673, Y676, M677, S678, S681 featured high cScore values and formed a cluster of coevolutionary centers in the S2 subunit (Figures 5,6). The multi-basic S1/S2 site in SARS-CoV-2 harbors multiple arginine residues and is involved in proteolytic cleavage of the S protein which is critical for viral entry into cells.^134^ A particular relevance of this site stems from the fact that sequence of the S1/S2 site enables cleavage by furin in SARS-CoV-2 but not in SARS-CoV or MERS-CoV viruses.^135^ The experimental data also showed that SARS-CoV S-mediated virus entry is based on sequential proteolytic cleavage at two distinct sites, with cleavage at the S1/S2 boundary (R667) promoting subsequent cleavage at the S2′ position (R797) triggering membrane fusion.^136^ Interestingly, our results indicated a high coevolutionary signal for R667 at the S1/S2 boundary but only a moderate Cscore for the conserved R797 position (Figure 5). These findings are generally consistent with coevolutionary patterns found in disordered protein regions showing that disordered residues whose function requires specific recognition and disorder-order transition upon binding can exhibit a high degree of coevolutionary signal.^137^

Although many coevolving centers in the S2 subunit are located inside the protein core and generally stable, these regions are involved in gigantic conformational rearrangements to the post-fusion state that require a nontrivial cooperation between these regions to dramatically rearrange the interaction network. Several important clusters of highly coevolving centers are localized in the HR1 regions (N925, A930, K947, N953, L959, F970, V976, L977, and L984) and CH regions (P987, E988, I993, D994, R995, I997, L1004, Y1007, T1027, and L1039) (Figures 5,6).

An interesting cluster of coevolving centers is formed by residues from different S regions surrounding the C-terminus of the central HR1-CH helices (Figure 6). This cluster included HR1 residues Q920, N925, F927, β-hairpin (BH) motif residue F1052, F898 (from connecting region 841-911) and several other hydrophobic positions F800 and F802 from the region upstream of the fusion peptide FP (816-SFIEDLLFNKVTLADAGF-833) (Figure 6B). These results showed that a number of the interface core and inter-protomer centers in the S2 subunit featured a significant coevolutionary signal. Consistent with the coevolutionary studies of protein complexes,^138^ we found that coevolutionary signal can be significant for the S2 positions involved in multiple interactions at critical junctures of UH, HR1 and CH regions. Hence, the increased structural and functional constraints for sites involved in significant number of inter-residue contacts can often imply the higher coevolution values. We found that both conservation and coevolutionary signals can increase for the S2 core residues involved in the inter-protomer interfacial contacts. However, the S2 core residues with the strongest coevolutionary signal and highest Cscore values could feature different level of conservation. In particular, the strongest coevolutionary centers in the S2 regions included fairly moderately conserved residues F1089, R983, N925 and L984 as well as strongly conserved A893, V915, L916, and A1190 positions. In general, our results indicated that S2 core residues subjected to more structural constraints and inter-residue contacts can exhibit the higher residue conservation and coevolution values.

To further probe the notion that the interface core residues can exhibit both the higher level of conservation and coevolution, we computed the average number of the inter-residue contacts for each S protein residue and aligned this distribution with the top Cscore positions in the S1/RBD and S2 regions (Figure 5C). The results indicated that coevolutionary centers tend to have a fairly significant number of the interacting contacts and can be involved in multiple interactions. In particular, coevolutionary hotspots in the RBD regions were often aligned with the peaks of the contact distribution, supporting the notion that the level of coevolution may be greater in residues involved in multiple interactions.^138^ It is worth noting, however, that sites with the largest number of inter-residue contacts may not necessarily correspond to the most conserved positions or residues with the highest CScore value. In fact, even though coevolutionary hotspots in the S2 subunit have a significant number of the inter-residue contacts, the distribution peaks corresponded to residues F718, V729, I742, F782 with moderate levels of conservation and coevolution (Figure 5C).

We also computed the distance probability distribution of coevolving directly coupled residue pairs in the studied SARS-CoV-2 S structures (Figure 5D). The profile showed several local maxima at 3.2 Å, 4.7 Å and a much broader area with a shallow peak near 7 Å - 8Å (Figure 5D). It is evident that the first two peaks reflect physical interactions between residues including hydrogen bonding and hydrophobic residue pairs. Hence, direct coevolutionary residue couplings in the SARS-CoV-2 S structures are dominated by spatially proximal residue pairs, that is consistent with large-scale investigations of direct coevolutionary couplings in proteins suggesting that coevolutionary signals are stronger for locally interacting residues than for residues involved in long-range interactions in allosteric networks.^139^ Nonetheless, the distribution also highlighted another intermediate range of coevolutionary couplings at 7 Å - 8Å that is beyond direct inter-residue physical contacts and may reflect strong couplings between spatially proximal functional regions (Figure 5D). This third distribution peak can correspond to coevolutionary couplings anchored by CTD1 regions (residues 529-591) in the S1 subunit that is believed to function as allosteric connector between RBD and FPPR regions by communicating signal from and to the fusion peptide.^26^

Hence, our results revealed the presence of a significant coevolutionary signal between functional regions separated by the medium-range distances which may help to facilitate a long-range cross-talk between distant allosteric regions in the S1 and S2 subunits.

Structural mapping of coevolutionary centers highlighted global connectivity of the coevolutionary network spanning from the epitope binding site towards the CTD1 region and regions in the S2 subunit (Figure 7). Collectively, these clusters could form modules of a coevolutionary network that may allow for efficient allosteric interactions and communications in the SARS-CoV-2 S proteins. In the SARS-CoV-2 complexes with H014, the contact interface is fairly large and involves a significant stretch of the RBD residues. Several high CScore residues W436, G413, F374, F377, and C379 are involved in the interactions with H014. In particular, multiple favorable interactions are formed by F377 with Y105, T58, S59, D60, Y50 of H014 and by C379 with N55, T58, G56,G57 positions of H014 (Figure 7A). S309 binding involves interactions with the highly conserved C336 and C361 positions that also correspond to coevolutionary hotspots and could anchor a network of evolutionary coupled residues in the S protein (Figure 7B). S2M11 interacts with the evolutionary conserved RBD sites F374 and W436 that also displayed high CScore values (Figure 7C). A smaller patch of coevolutionary centers is involved in contacts with S2E12 that connects the epitope with the S2 subunit via CTD1 region that serves as a mediating hub of coevolutionary clusters in the S1 (Figure 7D). Collectively, these clusters could form modules of a coevolutionary network that may allow for efficient allosteric interactions and communications in the SARS-CoV-2 S proteins.

**Figure 7.**
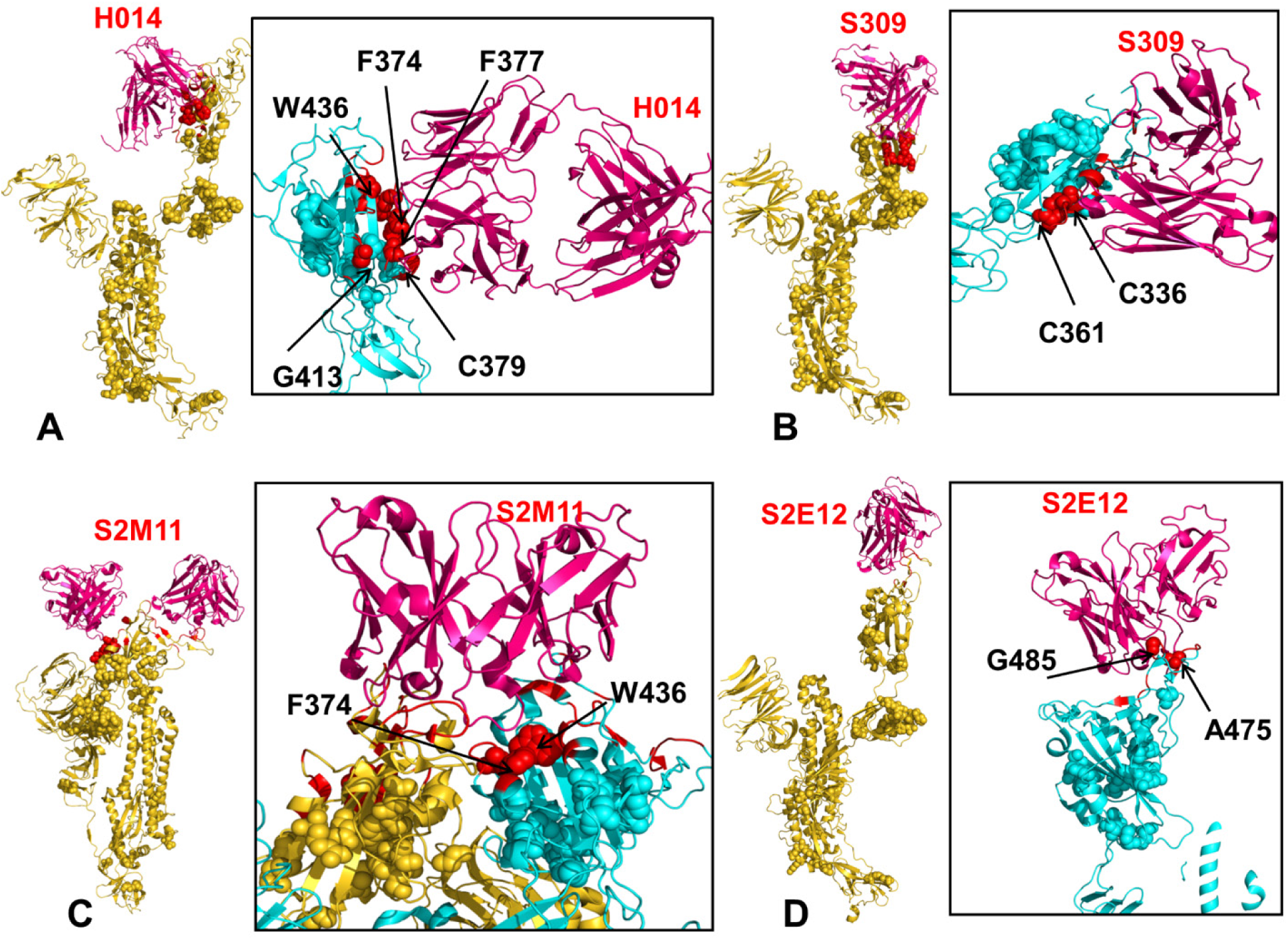
Structural maps of coevolutionary centers in the epitope regions of the SARS-CoV-2 complexes with Abs. (A) Structural map of coevolutionary centers in the S complex with H014 (pdb id 7CAI/7CAK) projected onto a single “up” protomer shown in green ribbons. The coevolutionary centers are in spheres and high CSscore hotspots from the binding epitope are in red spheres. A close-up of the H014 binding epitope with the coevolving centers involved in direct contacts with H014 in red spheres and annotated. (B) Structural map and close-up of coevolutionary centers in the S complex with S309 (pdb id 6WPT). The coevolving centers involved in direct contacts with S309 in red spheres and annotated. (C,D) Structural map and close-up of coevolutionary centers in the S complex with S2M11 (pdb id 7K43) and S2E12 (pdb id 7K4N). The coevolving centers involved in direct contacts with S2M11 and S2E12 are in red spheres and annotated.

### Conformational Dynamics and Mutational Scanning Reveal Modulation of Protein Stability and Binding Energy Hotspots of the SARS-CoV-2 Spike Complexes

We employed multiple CABS-CG simulations followed by atomistic reconstruction and refinement to provide a detailed comparative analysis of dynamic landscapes that are characteristic of the SARS-CoV-2 S trimer complexes with H014, S309, S2M11, and S2E12 Abs. Using these simulations, we examined how Ab binding could affect the global dynamic profiles of the closed, partially open, an open states revealing the important regions of flexibility (Figure 8). The analysis of the inter-residue contact maps (Figure S4, Supporting Information)^140^ and inter-residue distance maps (Figure S5, Supporting Information)^141^ in the SARS-CoV-2 S complexes with Abs indicated that the density of the interaction contacts is significantly greater in the densely packed S2 domains. The overall packing density of the closed S protein conformations complexed with S309 and S2M11 is also markedly higher as compared to the partially open and open states. Molecular simulations of the SARS-CoV-2 S complexes provided a quantitative picture of the differences in flexibility of the S protein states and the effect of Ab binding on modulation protein stability. A comparative analysis of the dynamics profiles showed that H014 binding can induce the significant dynamic changes by considerably reducing thermal fluctuations in the S1 regions of the Ab-interacting open protomers as compared to the unbound trimer form (Figure 8A,B). We observed small thermal fluctuations with RMSF < 1.0 Å for the S1 epitope positions (residues 368 to 386, 405 to 408 and 411 to 413, 439, and 503) that were considerably rigidified in both H014 complexes (Figure 8A,B). These findings are consistent with the experimental structural data suggesting that Ab-induced structural changes could trigger stabilization changes in both the RBD and NTD regions.^45^

**Figure 8.**
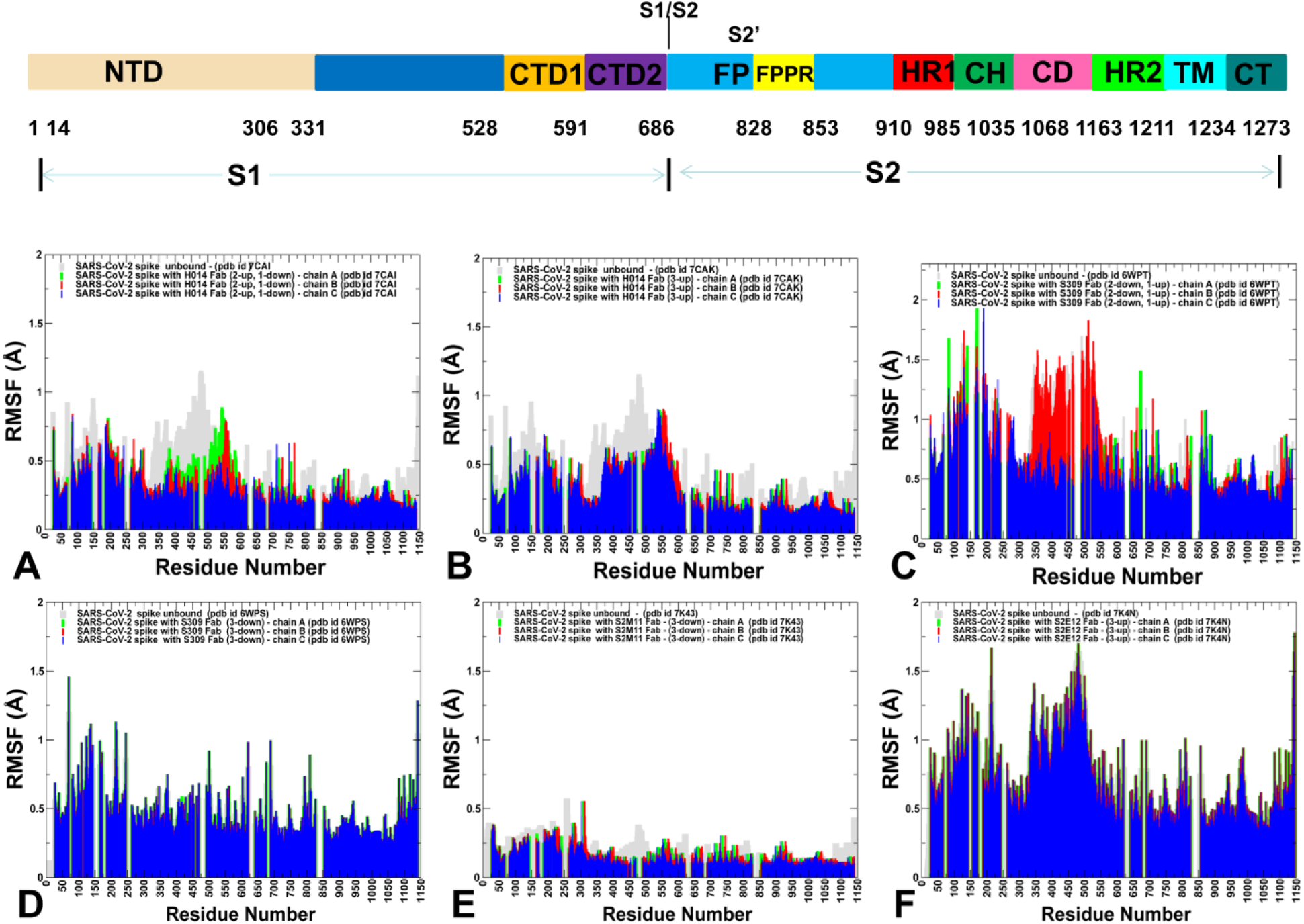
CABS-GG conformational dynamics of the SARS-CoV-2 S complexes. A schematic representation of domain organization and residue range for the full-length SARS-CoV-2 S protein is shown above conformational dynamics profiles. (A,B) The root mean square fluctuations (RMSF) profiles from simulations of the cryo-EM structures of the SARS-CoV-2 S trimer with H014. (C,D) The RMSF profiles from simulations of the cryo-EM structures of the SARS-CoV-2 S trimer with S309. (E) The RMSF profiles from simulations of the cryo-EM structures of the SARS-CoV-2 S trimer with all three RBDs in the closed form bound with S2M11. (F) The RMSF profiles from simulations of the cryo-EM structure of the SARS-CoV-2 S protein trimer with all three RBDs in the open-up form bound with S2E12. The profiles for protomer chains A,B and C are shown in green, red and blue bars respectively. The RMSF profiles for the unbound forms of S protein trimer are shown in light grey bars.

Conformational dynamics of the SARS-CoV-2 S protein complex with S309 showed only minor changes in the flexibility upon binding, particularly in the complex with S309 bound to 2 closed protomers (Figure 8C). In this case, the unbound open protomer displayed an appreciable flexibility, while the NTD regions of S309-bound closed protomers also showed some degree of mobility. In the S309 complex with 3 Abs bound to closed protomers, we found that stability of the closed S protein is protected, with only minor changes in the local dynamics between unbound and S309-bound S forms (Figure 8D). S2M11 functions by locking down the SARS-CoV-2 S trimer in the closed state through binding to a quaternary epitope. Conformational dynamics profile of the S protein complex with S2M11 in the closed form reflected this mechanism by featuring an extremely stable SARS-CoV-2 S conformation in which both S1 and S2 regions were virtually immobilized and displayed only very minor thermal fluctuations (Figure 8E). Interestingly, according to our analysis, this is the most stable bound form of the SARS-CoV-2 S protein among studied systems, suggesting that ultra-potent neutralization effect may be partly determined by the extreme thermodynamic stabilization of the closed-down S protein form. The mechanism of S2E12 neutralization of SARS-CoV-2 S protein is based on direct targeting of the RBM regions and interfering with ACE2 binding. A relatively small binding epitope in the S2E12 complex with the fully open form of S protein produced the dynamics profile where NTD and RBD regions showed an appreciable degree of mobility, while the S2 regions were mostly immobilized (Figure 8F). The important finding of this analysis was that H014, S309 and S2M11 Abs can exert modulation of the conformational dynamics leading to a significant stabilization of both S1 and S2 regions in the open protein forms, which may effectively counteract the intrinsic flexibility of the receptor-accessible, open S conformations and thus induce potent neutralization effects.

To establish connection between dynamics and energetics of the SARS-CoV-2 binding, we employed the conformational ensembles generated in simulations and performed a systematic alanine scanning of the protein residues (Figure 9). The results revealed a wide range of important binding hotspots in the S protein complexes with H014 (Figure 9A,B). This is consistent with the dynamics profile showing a broad stabilization of the RBD regions, including the epitope residues and RBM positions. In particular, alanine scanning showed a significant contribution of conserved RBD residues F374, F377, K378, C379, Y380, P384, T385 as well as N437, V503, Y508 (Figure 9B). Among binding energy hotspots we detected some of the highly conserved positions and several coevolutionary centers such as F374, F377, and C379 residues. We argue that through interactions with major coevolutionary centers in the conserved RBD epitope, H014 may exert its long-range effect by propagating binding signal through clusters of proximal coevolutionary pairs in the RBD and CTD1 regions. The noticeably fewer number of binding hotspots were seen in the S309 complexes with partially closed (2-down) and fully closed forms of the S protein (Figure 9C,D). The determined binding hotspots L335, P337, T345, and L441 are characterized by only moderate conservation and CScore values. S309 also makes weaker contacts with conserved and coevolutionary important RBD centers C336 and C361. However, the binding free energy changes caused by alanine mutations in these positions are fairly moderate (∼0.7 - 0.8 kcal/mol). The binding energy hotspots in the S2M11 complex with S protein occupy two different regions, where one group includes conserved RBD sites F374 and W436 that also displayed high CScore values (Figure 9E). Another group of binding energy hotspot positions includes moderately conserved residues Y449, F456, F484, F486, Y489 that form a critical patch of the RBM binding interface with the host receptor.

**Figure 9.**
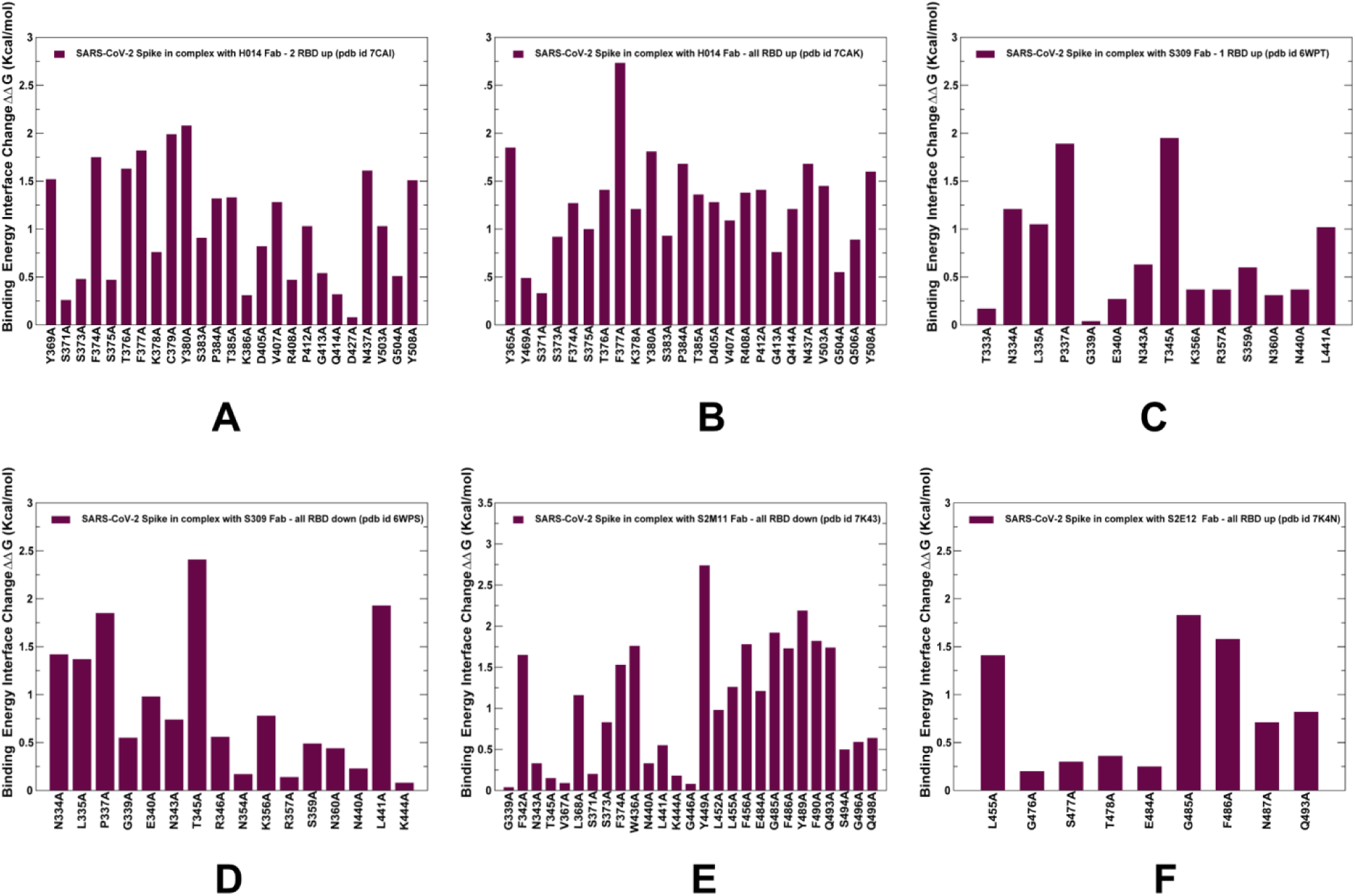
Alanine scanning of the binding epitope residues in the SARS-CoV-2 S complexes with a panel of Abs. The binding free energy changes upon alanine mutations for the epitope residues in the SARS-CoV-2 S complex with H014 - two RBDs in the open state, pdb id 7CAI (panel A), SARS-CoV-2 S complex with H014 - three RBD in the open state, pdb id 7CAK (panel B), SARS-CoV-2 S complex with S309 - two RBDs in the closed form, pdb id 6WPT (panel C), SARS-CoV-2 S complex with S3090 - three RBDs in the closed form, pdb id 6WPS (panel D), SARS-CoV-2 S complex with S2M11 - three RBDs in the closed form, pdb id 7K43 (panel E), SARS-CoV-2 S complex with S2E12 - three RBDs in the closed form, pdb id 7K4N (panel F). The computed binding free energy changes values are shown in bars. The binding interface residues are determined for each complex based on the average interaction contacts that persist during simulation of a given complex.

We previously showed that a conserved segment 486-FNCYFPL-492 in the RBD region emerged as a central binding energy hotspot in the SARS-CoV-2 complex with ACE2 receptor.^62^ Hence, through binding to two adjacent protomers S2M11 can simultaneously block interface with ACE2 and influence long-range couplings using a network of coevolutionary coupled residues. In general, the results indicated that the interactions S014 and S2M11 Abs can lead to stabilization of the S conformations and emergence of multiple binding energy hotspots. By targeting these centers H014 and S309 can exert their neutralizing effect by achieving a strong binding affinity with the SARS-CoV-2 S protein and also by strengthening long-range couplings of S1 an S2 regions (Figure 9).

### Hierarchical Network Modeling Reveals Mediating Centers of Allosteric Interactions in the SARS-CoV-2 Spike Complexes

We applied a hierarchical-based network modeling approach in which the residue interactions and network couplings are described with the increasing level of atomistic details and complexity. First, a protein contact network was implemented to highlight the topological role of residues in protein structure activity and identify residues mostly responsible for signal transmission throughout the protein structure. In this simplified model, the protein residues correspond to network nodes and inter-residue contacts are considered as active links based on distance criteria as described in our previous studies.^98–100^ Based on the hierarchical clustering algorithm, we computed the average participation coefficient *P* values that measure the contribution of residue nodes in communication between different clusters (functional domains). To focus analysis on several prominent cases, we reported the communicating residues in the SARS-CoV-2 structures bound with H014 (Tables S7,S8, Supporting Information) and S309 (Tables S9,S10, Supporting Information). The results indicated that the majority of the inter-cluster communcating sites are localized in the RBD and especially CTD1 regions for SARS-CoV-2 S complexes with H014 (Tables S7,S8, Supporting Information). In this case, by attentuating mobility of the interacting RBD regions H014 binding may activate allosteric interaction networks and communications between S1 and S2 regions with CTD1 residues acting as global mediating centers of long-range interactions. The distribution of communicating positions in the SARS-CoV-2 S complexes with S309 (Tables S9,S10, Supporting Information) revealed an appreciably larger number of potential mediatig centers with significant communication propensities. Moreover, these positions corresponded to different regions, including a significant number of mediating hubs in the UH, CH and HR1 regions of S2 subunit as well as residues in the CTD1 regions of S1. These preliminary findings suggested that allosteric interaction networks in the SARS-CoV-2 S complexes with S309 could be broadly distributed, which can arguably reflect strengthening of allosteric couplings between S1 and S2 subunits as S309 locks the down-regulated form of the S protein. In the framework of the hierarchical approach, we also explored a more detailed model of the residue interaction networks by using a graph-based representation with residues as network nodes and the inter-residue edges defined by both dynamic correlations^104^ and coevolutionary residue couplings ^105^ as detailed in our previous studies.^105–107^ Using the results of simulations, the ensemble-averaged distributions of the betweenness centrality were computed for the SARS-CoV-2 S complexes with Abs (Figure 10). We found that the high centrality residues can be assembled in tight interaction clusters localized in the key functional regions of the S protein. In the SARS-CoV-2 S protein complexes with H014, the centrality profiles featured strong and dense peaks in the RBD and CTD1 regions of S1 as well as another peak in the CH region of S2 (residues 986-1035) (Figure 10A,B). The centrality peaks also aligned well with the hinge centers of S1 (residues 315-320, 569-572), indicating that these dynamically important control points could also mediate communication in the residue interaction networks. The network centrality analysis also revealed clusters of distribution peaks in the SARS-CoV-2 S complexes featuring the fully closed conformation (Figure 10D,E). In these structures S309 and S2M11 induce a strong stabilization effect and lock the S protein in the closed state. According to our results, these structurally stable states can also feature a broadly distributed allosteric network mediated by functional sits in both S1 and S2 subunits, primarily CTD1 regions (residues 529-591) UH (residues 736-781), CH (residues 986-1035), and β-hairpin (BH) region (residues 1035-1071). The dominant clusters of centrality peaks located in the RBD and CTD1 regions of S1 and CH regions of S2 can be seen in the S complex with S2E12 (Figure 10F). This showed that S2E12 binding may activate the increased mediating capacity of CTD1 regions and strengthen allosteric interactions between S1 and S2 regions.

**Figure 10.**
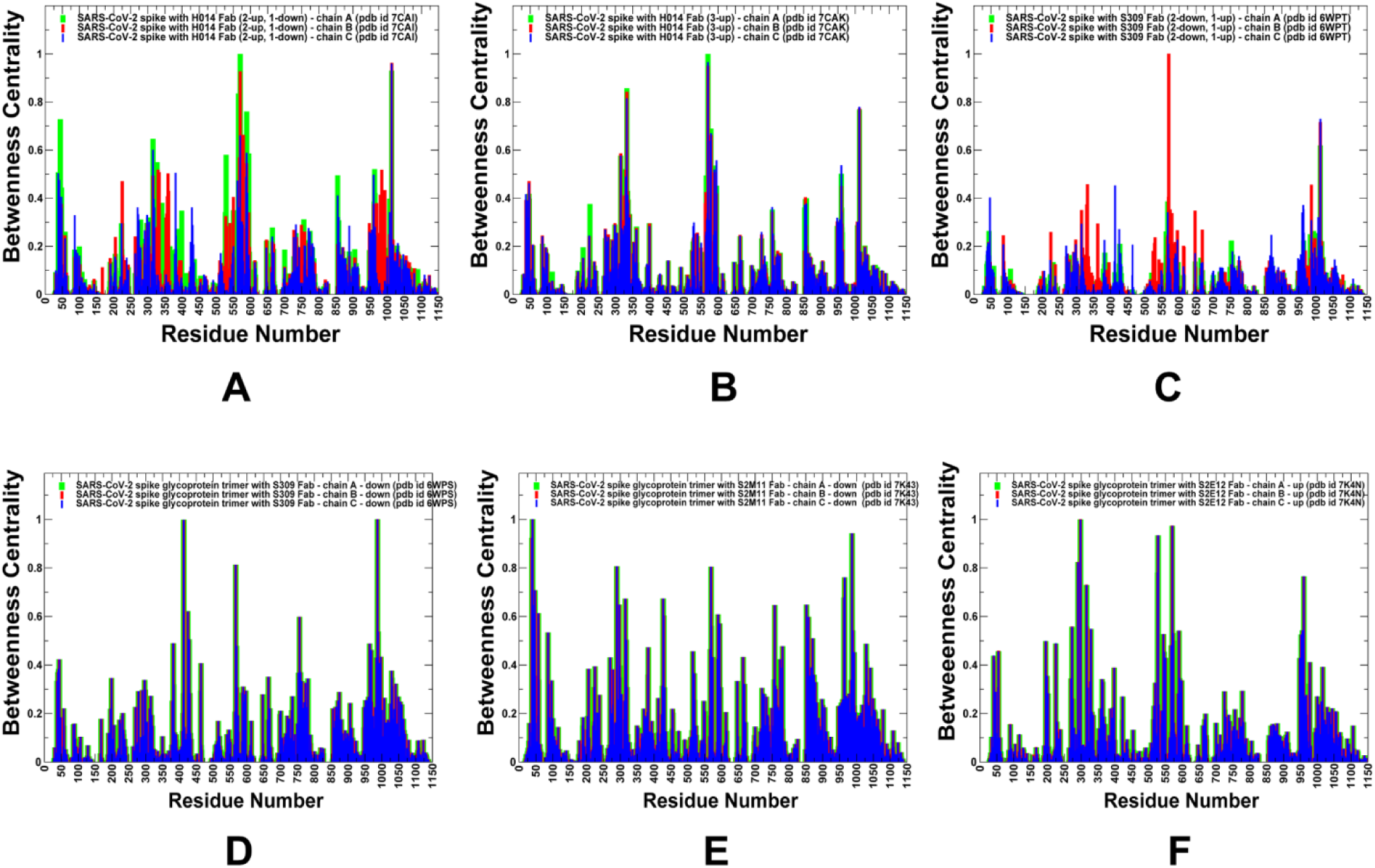
The residue-based betwenness centrality profiles in the SARS-CoV-2 S complexes with a panel of Abs. The centrality values are computed by averaging the results over 1,000 representative samples from CABS-CG simulations and atomistic reconstruction of trajectories. (A) The centrality profile is shown for the SARS-CoV-2 S complex with H014 - two RBDs in the open state (A), SARS-CoV-2 S complex with H014 - three RBD in the open state (B), SARS-CoV-2 S complex with S309 - two RBDs in the closed form (C), SARS-CoV-2 S complex with S309 - three RBDs in the closed form (D), SARS-CoV-2 S complex with S2M11 - three RBDs in the closed form (E), SARS-CoV-2 S complex with S2E12 - three RBDs in the closed form (F). The profiles for protomer chains A,B and C are shown in green, red and blue bars respectively.

Structural mapping of high centrality sites highlighted differences between network organizations in the SARS-CoV-2 complexes (Figure 11). In the complexes with H014 the high centrality sites are concentrated near CTD1 regions that could strengthen couplings at the inter-domain boundaries between S1 and S2 (Figure 11A,B). We argue that H014 binding may increase the allosteric potential of the RBD and CTD1 regions and activate communication between the RBD and S2 via CTD1 regions. Of particular interest is a dense network of mediating centers in the complexes with S309 and S2M11 (Figure 11C-E), showing that these Abs may facilitate a broad allosteric interaction network between S1 and S2 functional regions.

**Figure 11.**
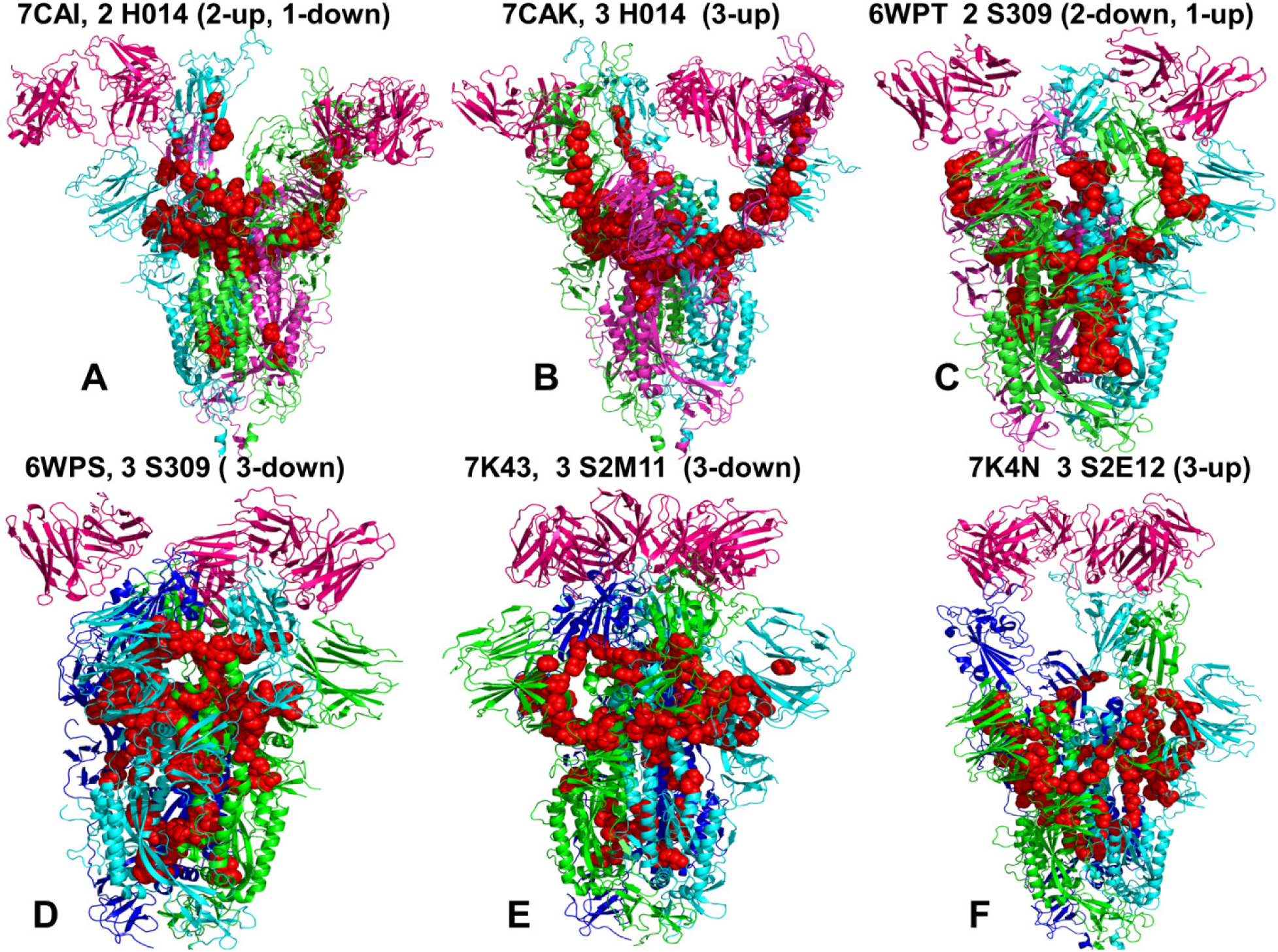
Structural maps of high centrality clusters in the SARS-CoV-2 S complexes. (A) Structural map for the SARS-CoV-2 S complex with H014 - two RBDs in the open state (A), SARS-CoV-2 S complex with H014 - three RBD in the open state (B), SARS-CoV-2 S complex with S309 - two RBDs in the closed form (C), SARS-CoV-2 S complex with S3090 - three RBDs in the closed form (D), SARS-CoV-2 S complex with S2M11 - three RBDs in the closed form (E), SARS-CoV-2 S complex with S2E12 - three RBDs in the closed form (F). The protomer A is shown in green ribbons, protomer B in cyan ribbons, and protomer C in blue ribbons. The bound Abs are depicted in dark pink-colored ribbons. The high centrality residue clusters are shown in red spheres.

### Perturbation Response Scanning Identifies Regulatory Hotspots of Allosteric Interactions in Different Conformational States of the SARS-CoV-2 Spike Trimer

Using the PRS method^112, 113^ we quantified the allosteric effect of each residue in the SARS-CoV-2 complexes in response to external perturbations. PRS analysis produced the residue-based effector response profiles in different functional states of the unbound SARS-CoV-2 S trimer (Figure 12) and SARS-CoV-2 S complexes with H014, S309, S2M11, and S2E12 Abs (Figure 13). The effector profiles estimate the propensities of a given residue to influence dynamic changes in other residues and are applied to identify regulatory hotspots of allosteric interactions as the local maxima along the profile. The central hypothesis tested in the PRS analysis is that Ab binding can incur measurable and functionally relevant changes by modulating the effector profiles of the unbound SARS-CoV-2 S protein timer. Moreover, we conjectured that binding can differentially affect the effector response profiles and allosteric interaction networks in distinct functional forms of the SARS-CoV-2 S protein. By systematically comparing the PRS profiles in the unbound and bound S protein forms, we determined the distribution of regulatory allosteric centers and clarified the role of specific functional regions in controlling allosteric conformational changes.

**Figure 12.**
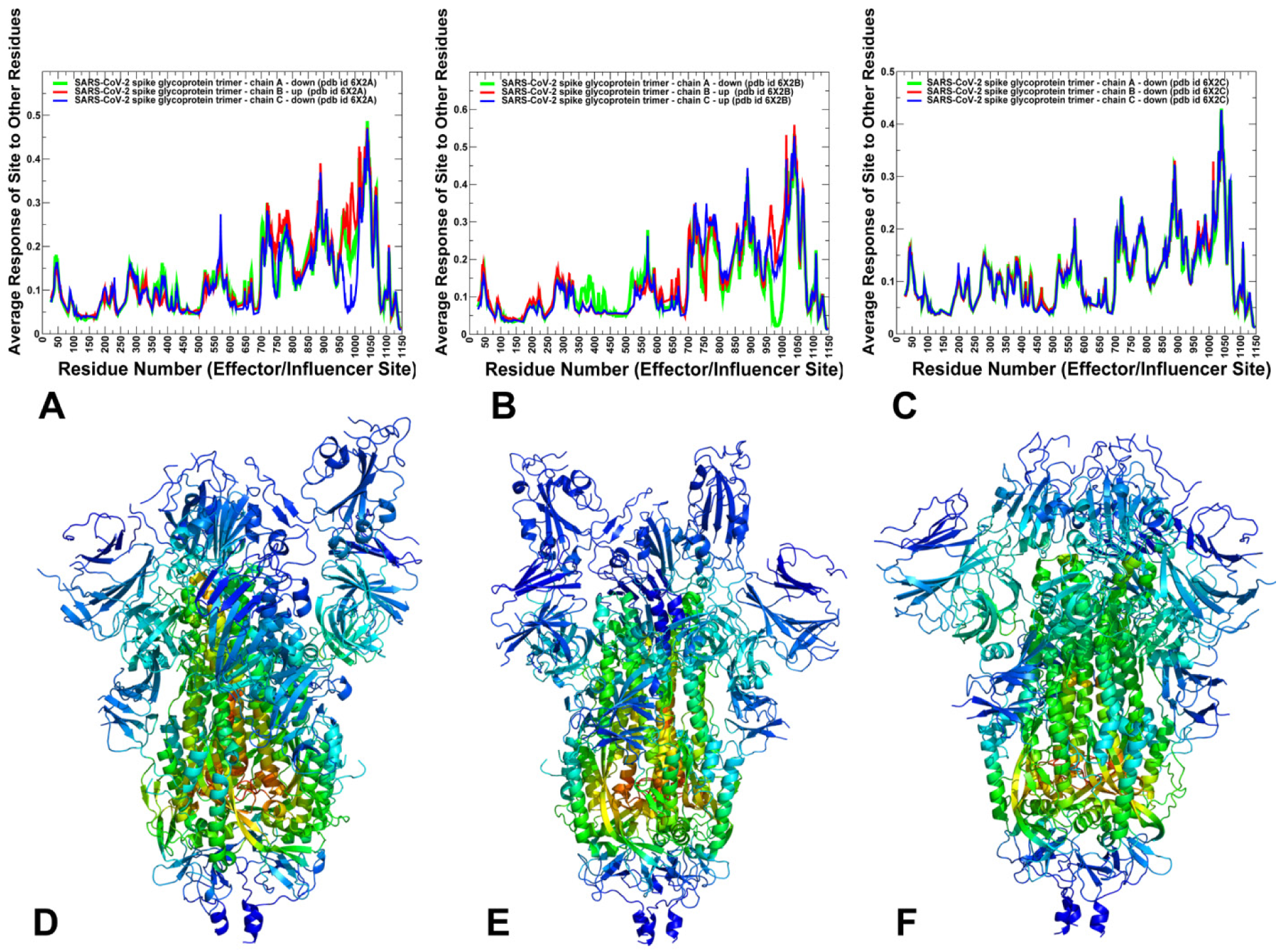
The PRS effector profiles in the closed, partially open and open states of the SARS- CoV-2 spike trimers. (A) The PRS effector profile is shown for the ligand-free SARS-CoV-2 S trimer in the partially open (1 RBD up) conformation (pdb id 6X2A). (B) The PRS effector profile for the ligand-free SARS-CoV-2 S trimer in the open (2 RBDs up) conformation (pdb id 6X2B). (C) The PRS effector profile for the ligand-free SARS-CoV-2 S trimer in the closed (3 RBDs-down) prefusion conformation (pdb id 6X2C). The profiles are shown for protomer in green lines, protomer B in red lines, and protomer C in blue lines. Structural maps of the PRS effector profiles are shown for the partially open state of the SARS-CoV-2 S prefusion trimer (D), open state (E), and closed state (F). The color gradient from blue to red indicates the increasing effector propensities.

**Figure 13.**
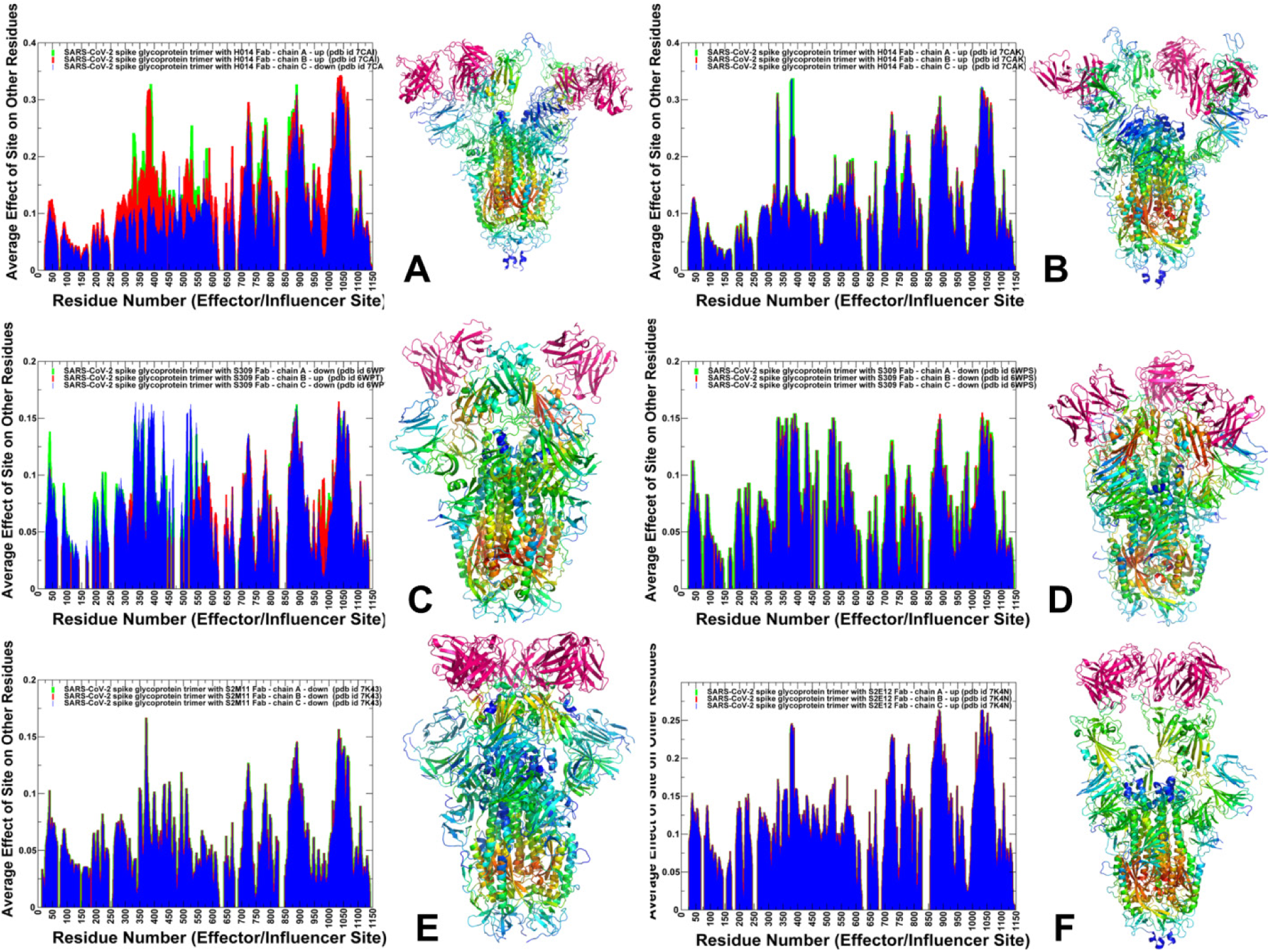
The PRS effector profiles for the SARS-CoV-2 S complexes with Abs. (A) The PRS effector distributions and structural maps of the effector profile are shown for the SARS-CoV-2 S complex with H014 - two RBDs in the open state (A), SARS-CoV-2 S complex with H014 - three RBD in the open state (B), SARS-CoV-2 S complex with S309 - two RBDs in the closed form (C), SARS-CoV-2 S complex with S3090 - three RBDs in the closed form (D), SARS-CoV-2 S complex with S2M11 - three RBDs in the closed form (E), SARS-CoV-2 S complex with S2E12 - three RBDs in the closed form (F). The profiles are shown for protomer in green bars, protomer B in red bars, and protomer C in blue bars. Structural maps of the PRS effector profiles are shown with the color gradient from blue to red indicating the increasing effector propensities.

To establish the baseline for comparison of allosteric profiles in the SARS-CoV-2 complexes we first computed PRS effector profiles for the unbound S protein in the closed (3 RBDs-down), partially open (2RBDs-down, 1RBD-up) and open states (1RBD-down, 2 RBDs-up) (Figure 12). In this analysis of the unbound forms of the SARS-CoV-2 S trimer, we used the cryo-EM structure of the full-length quadruple mutant (A507L T572I F855Y N856I) in the all-down closed prefusion conformation (pdb id 6X2C), the partially open (1-up) conformation (pdb id 6X2A), and the open (2-up) conformation (pdb id 6X2B).^28^ These structures revealed thermodynamic and dynamic differences between different states in which dynamic switch centers are responsible for modulation of allosteric changes between the closed and open S states.^28^ By using these structures as reference states for the PRS analysis, we also examined how Ab binding can alter the localization and effect of these regulatory switch points on allosteric interactions.

The results showed that the effector profile in the unbound S structures can remain largely conserved, displaying the largest peaks in the functional regions of the S2 subunit (Figure 12A-C). The major effector peaks corresponded to residues 756-758 in the UH region, HR1 region (residues 910-985), residues 886-890 and BH region (residues 1035-1071). In the closed and partially open forms, we also observed a secondary peak corresponding to CTD1 regions (residues 529-591) in the S1 subunit. Only a minor effector density was seen in the RBD regions (Figure 12A-C). At the same time, these regions can serve as primary sensors of allosteric signaling in the S trimers for the partially open and open states (Figure S6, Supporting Information). These results are consistent with our latest studies of the SARS-CoV-2 S structures demonstrating that the broadly distributed effector density can be seen only in the fully locked closed state, while allosteric couplings in more dynamic closed and open forms may be largely governed by the regulatory centers in core of the S2 subunit.^64^ The reduced density of the effector centers in the RBD regions indicated that the allosteric signaling in the dynamic closed and open forms may be primarily one-directional, in which allosteric centers in the S2 core regions could dictate allosteric changes in the RBD regions. Based on this preliminary evidence, we suggested that efficient allosteric signaling between S1 and S2 subunits and broad allosteric networks may be salient features of the thermodynamically locked closed S form.

We conjectured that Ab binding at different RBD epitopes may affect not only local interactions and stability near the binding sites but also have long-range effect by modulating allosteric effector potential of SARS-CoV-2 S regions and altering allosteric interaction networks. This can allow for highly cooperative motions in which many spatially distributed effector residues are in the allosteric network that links distant S1 and S2 functional regions. In this model, allostery requires an effector ligand to stabilize the interactions in the closed S state over those in the open S states.

The central result of this analysis is that the studied neutralizing Abs could effectively restore allosteric potential of the RBD and CTD1 regions in the closed and open states without compromising the effector potential of S2 regions, thereby introducing ligand-induced cooperativity and strengthening the broad allosteric interaction network (Figure 13). Indeed, the results showed that H014 binding induced significant changes in the effector profile of the RBD and CTD1 regions for “up” protomers while preserving and enhancing the effector capacity of the CH and CD residues (Figure 13A,B). In the partially open form of the SARS-CoV-2 complex, the effector peak center corresponded to RBD residues S383, F377, K378, C379, Y380, G381, V382, S383, P384 involved in direct productive interactions with H014 (Figure 13A). Several of these effector centers (F377, C379, and S383) also corresponded to conserved positions implicated in mediating coevolutionary couplings. In addition, we noticed a significant increase of the effector potential for the CTD1 regions in the open protomers. Similar findings were observed in the open form (3RBDs - up) of the SARS-CoV-2 S complex with H014 where the allosteric potential of RBD regions was markedly enhanced (Figure 13B). Structural maps of the effector profiles further illustrated these findings by showing the increased effector density penetrating into the RBDs of protomers that interact with H014 (Figure 13A,B).

These results suggested that H014 binding could modulate the effector propensities of the RBD residues and restore the allosteric potential of the S1 regions, leading to strengthening of the network of allosteric interactions in the complex. The effector profiles also indicated the increased density and clustering of effector peaks distributed across RBD, CTD1, UH and CH regions (Figures 13A,B).

The effector profiles of SARS-CoV-2 complexes with S309 showed the significantly increased effector potential of the RBD and CTD1 regions, which is manifested in the emergence of dominant and broad peaks in these S1 regions (Figure 13C,D). Among emerging effector peaks in the RBD regions are T333, N334, C361, V362, A363, L390 and C391. These residues correspond to highly conserved and structurally stable positions involved in stabilizing disulfide linkages, C336–C361 and C391–C525 that anchor the β-sheet structure in the RBD regions. The S309-induced modulation of allosteric effector propensities could be also amplified by the fact that this Ab appears to thermodynamically stabilize partially closed and closed forms of the S protein where two or all three RBDs assume the down-regulated conformation (Figure 13C,D). Structural maps highlighted a significant expansion of the effector density towards S1 regions and boundaries between S1 and S2 subunits (Figure 13C,D). By strengthening allosteric couplings between S1 and S2 subunits, S309 could arguably lock the down-regulated form to ensure S1- based protection of the fusion machinery.

S2M11 binds a quaternary epitope comprising distinct regions of two neighboring RBDs within an S trimer and induced stabilization of SARS-CoV-2 S in the closed conformational state. We found that S2M11 binding can promote the increased effector potential of conserved and structurally stable residues that are not directly involved in binding contacts (Figure 13E). Indeed, S2M11 can induce the increased effector potential in the RBD regions, particularly residues F374, F377, K378 C379, Y380 (Figure 13E). S2E12 recognizes an RBD epitope overlapping with the RBM that is partially buried at the interface between protomers in the closed S trimer and therefore S2E12 can only interact with open RBDs. According to our results, S2E12 binding can cause a similar redistribution of allosteric effector potential in the S1 regions and activate effector capacity of conserved stretch of residues in the β-sheet of the RBD regions (Figure 13F). Strikingly, S2E12-induced modulation of the effector propensities in the fully open S conformation can effectively restore allosteric potential for the RBD and CTD1 S1 regions, while strengthening peaks in the UH and CH regions of S2 subunit. As a result, as highlighted by structural projection of the effector profiles, the potential effector centers become spatially distributed across the S trimer structure (Figure 13F).

The PRS sensor profiles describe the propensity of flexible residues to serve as carriers of large allosteric conformational changes. We found that Ab binding can induce changes in the shape and peaks of sensor profiles in the partially open and closed S forms (Figure S7, Supporting Information). The sensor peaks in the RBD regions that are prevalent in the unbound S protein become partially suppressed, featuring now a broader distribution of small peaks localized across S1 and S2 regions. This suggested that the sensor propensities of the RBD regions can be significantly reduced in the SARS-CoV-2 S complexes, which reflects conformational and dynamic constraints imposed by Abs on large structural changes.

The results of the PRS analysis also suggested that allosteric mechanisms underlying Ab binding to S proteins may bear signs of ligand-induced cooperativity in which the effector can shift the distribution of local interactions and energies for many residues.^142^ Based on our results, we argue that Abs can induce a switch from a moderately cooperative population-shift mechanism of the unbound S protein to a highly cooperative ligand-induced allosteric mechanism. While the allosteric interaction network of the unbound S protein in the population-shift mechanism tends to be more dispersed and smaller, binding can induce a large and dense allosteric network that efficiently couples local changes in the distant S1 and S2 regions.

In this context, it is worth noting that cooperative allosteric mechanisms with a broad allosteric network tend to better withstand mutations in the functional regions without significant deleterious consequences for protein function.^142^ Accordingly, it may be suggested that the ligand-induced cooperative allosteric effect produced by Ab binding may enhance resistance against mutations so that mutational changes would not easily alter conformational preferences and expose the RBD regions to interactions with the host receptor.^143–147^ In some contrast, a less cooperative population-shift mechanism in the unbound S protein may be more susceptible and vulnerable to mutations of residues in the communication network, which may allow individual mutations at the regulatory switch centers to alter conformational equilibrium and potentially increase population of the receptor-accessible open S conformations.

## Conclusions

This study examined molecular mechanisms underlying SARS-CoV-2 S protein binding with a panel of highly potent Abs through the lens of coevolutionary relationships and ligand-induced modulation of allosteric interaction networks. The results revealed key functional regions and regulatory centers that govern coevolutionary couplings and allosteric interactions in the SARS-CoV-2S protein complexes. We found that Ab-specific targeting of coevolutionary hotspots in the S protein can allow for efficient modulation of long-range interactions between S1 and S2 units by propagating signal through clusters of spatially proximal coevolutionary coupled residues. The results revealed strong coevolutionary signal between functional regions separated by the medium-range distances which may help to facilitate a long-range cross-talk between distant allosteric regions. Conformational dynamics and binding energetics analyses showed that binding of Abs can lead to significant stabilization of both S1 and S2 regions which may be relevant in rationalization of potent neutralization effects. The PRS analysis of the unbound and bound SARS-CoV-2 S proteins showed that Abs can promote formation of highly cooperative and broad allosteric networks that restore and enhance couplings between S1 and S2 regions, thereby inhibiting dissociation of S1 subunit from the spike apparatus required for membrane fusion. By systematically comparing the PRS profiles, we clarified the role of specific functional regions in regulating allosteric interactions. The results of this study provide a novel insight into allosteric regulatory mechanisms of SARS-CoV-2 S proteins showing that Abs can uniquely modulate signal communication providing a plausible strategy for therapeutic intervention by targeting specific hotspots of allosteric interactions in the SARS-CoV-2 proteins.

## AUTHOR INFORMATION

The authors declare no competing financial interest.

## Acknowledgment

This work was partly supported by institutional funding from Chapman University. The author acknowledges support by the Kay Family Foundation Grant A20-0032.

## ABBREVIATIONS

SARS: Severe Acute Respiratory Syndrome
RBD: Receptor Binding Domain
ACE2: Angiotensin-Converting Enzyme 2 (ACE2)
NTD: N-terminal domain
RBD: receptor-binding domain
CTD1: C-terminal domain 1
CTD2: C-terminal domain 2
FP: fusion peptide
FPPR: fusion peptide proximal region
HR1: heptad repeat 1
CH: central helix region
CD: connector domain
HR2: heptad repeat 2
TM: transmembrane anchor
CT: cytoplasmic tail

## Notes

### Competing Interest Statement

The authors have declared no competing interest.

