## Supplemental Information for "Coevolutionary Analysis and Perturbation-Based Network Modeling of the SARS-CoV-2 Spike Protein Complexes with Antibodies: Binding-Induced Control of Dynamics, Allosteric Interactions and Signaling"

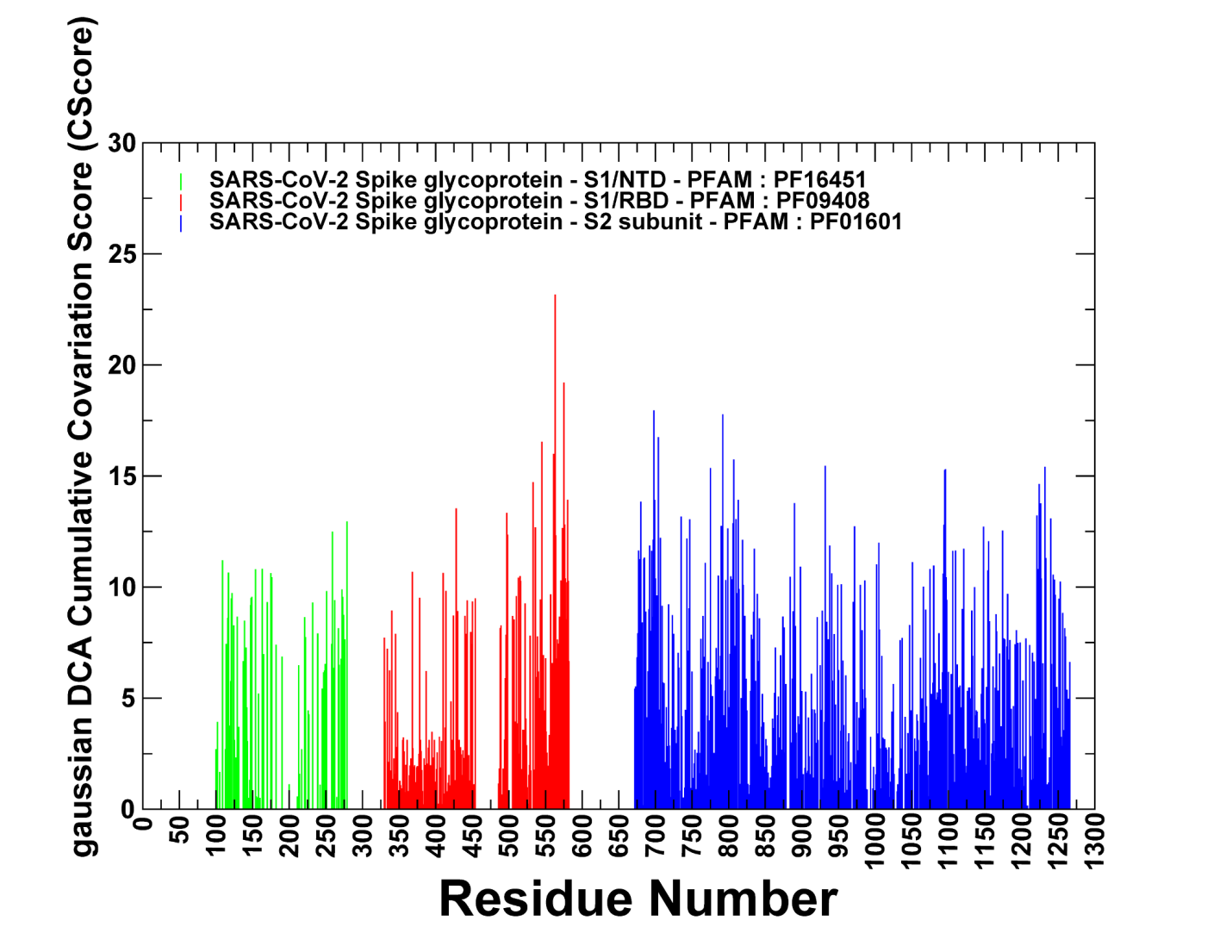
**Figure S1.** Coevolutionary profiles of the SARS CoV-2 S proteins. (A) The Gaussian DCA-based coevolutionary Cscore profile for the SARS-CoV-2 S proteins (P0DTC2: SPIKE_SARS2 sequence numbering). The Cscore values are shown for the S1-NTD residues in green bars (Pfam:PF16451), for the RBD in red bars (Pfam:PF09408) and for S2 regions in blue bars (Pfam:PF01601). Three Pfam domains were utilized corresponding to S1, the NTD (N-terminal, Pfam:PF16451, Uniprot P0DTC2: SPIKE_SARS2 sequence numbering), the RBD (Pfam:PF09408, Uniprot P0DTC2: SPIKE_SARS2 sequence numbering) and the new C-terminal domain, CTD (Pfam:PF19209, Uniprot P0DTC2: SPIKE_SARS2 sequence numbering). S2 is described in the family Pfam:PF01601 (Uniprot P0DTC2: SPIKE_SARS2 sequence numbering).

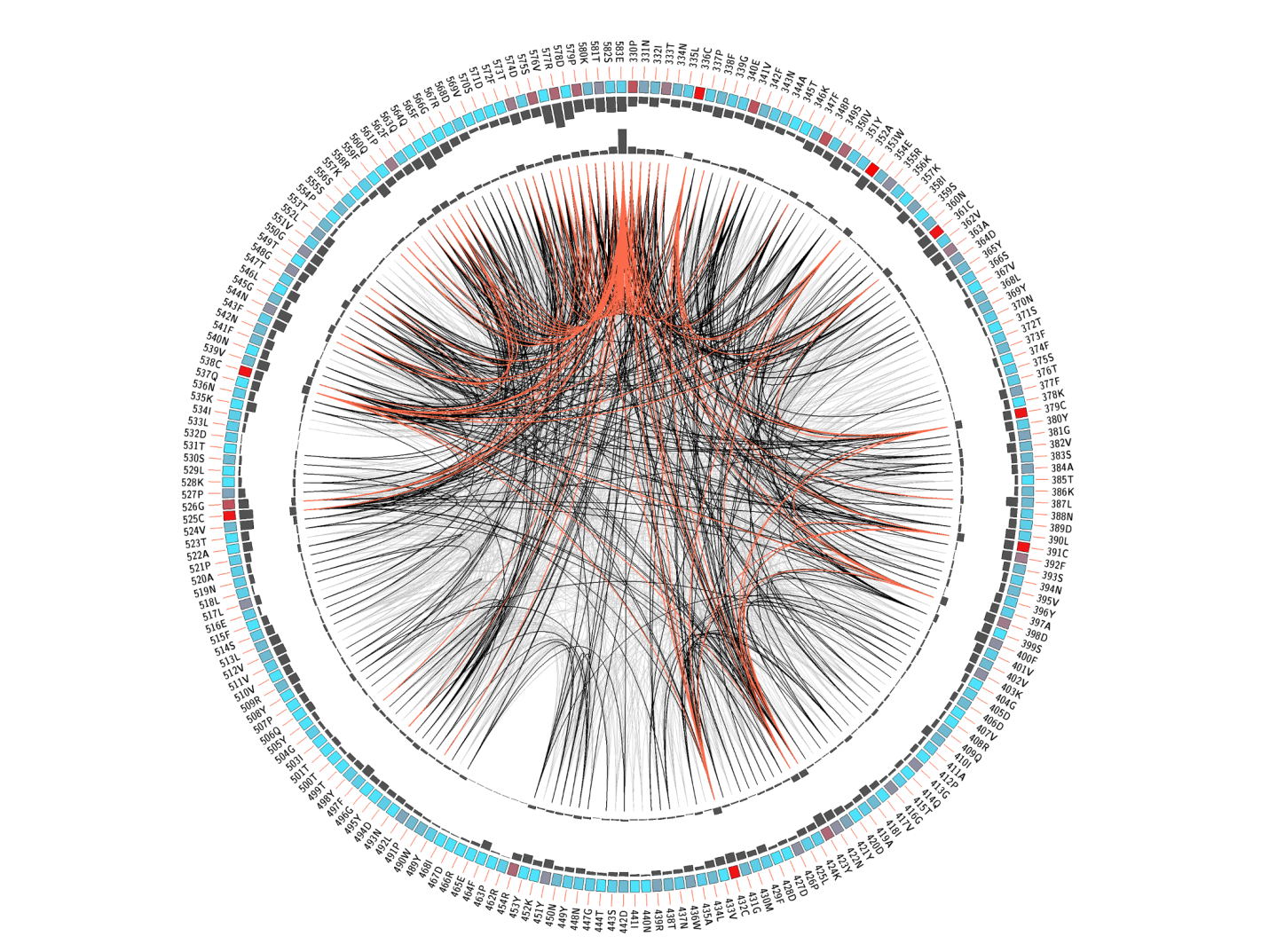
**Figure S2.** The network connectivity of coevolving residue pairs in S1 regions of the SARS-CoV-2 S protein is shown by a sequential circular representation. S2 is described in the family Pfam:PF01601. The labels in the first (outer) circle indicate the alignment position and the amino acid code of the reference sequence. The colored square boxes of the second circle indicate the MSA position conservation (highly conserved positions are in red, while less conserved ones are in blue).The fourth circle shows the CSscore values as histograms, facing outwards. Each node represents a position in the MSA and lines between nodes in the circle connect pairs of coevolving positions. Node color represents site conservation: red for highly conserved sites and blue for less conserved sites.

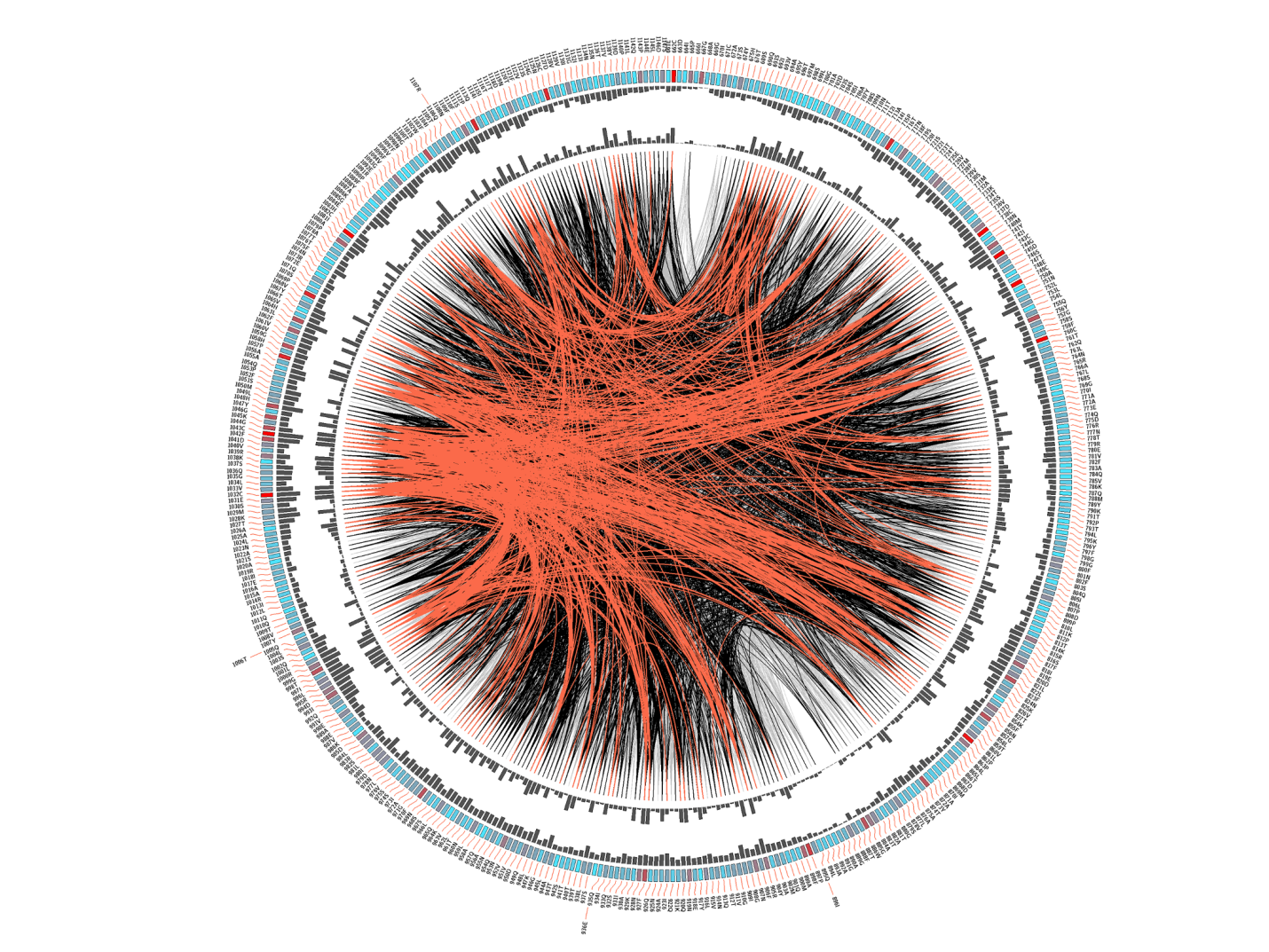
**Figure S3.** The network connectivity of coevolving residue pairs in S2 regions of the SARS-CoV-2 S protein is shown by a sequential circular representation. S2 is described in the family Pfam:PF01601. The labels in the first (outer) circle indicate the alignment position and the amino acid code of the reference sequence. The colored square boxes of the second circle indicate the MSA position conservation (highly conserved positions are in red, while less conserved ones are in blue).The fourth circle shows the CSscore values as histograms, facing outwards. Each node represents a position in the MSA and lines between nodes in the circle connect pairs of coevolving positions. Node color represents site conservation: red for highly conserved sites and blue for less conserved sites.

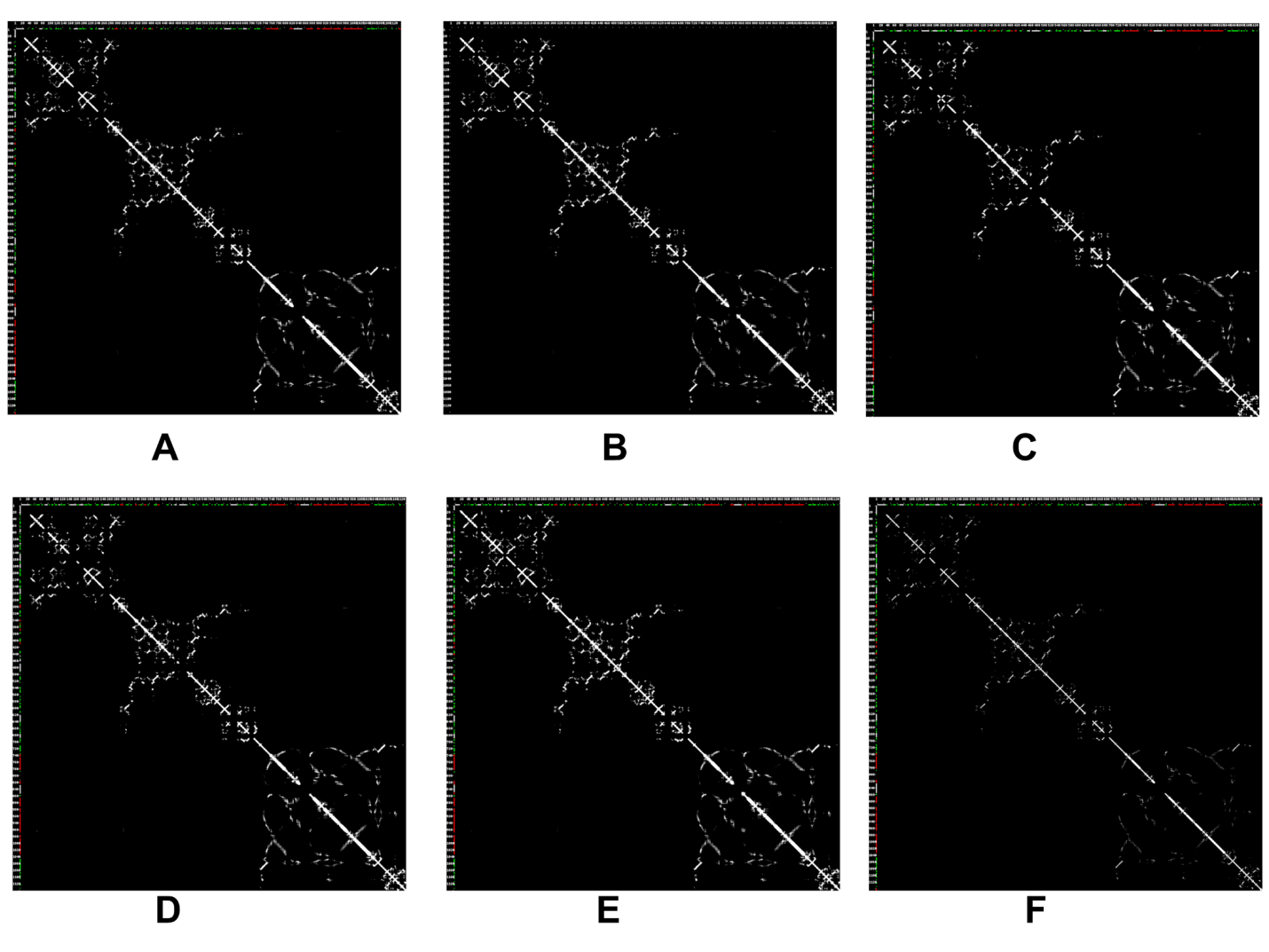

**Figure S4.** The inter-residue contact maps the SARS-CoV-2 S complexes with Abs. (A) The inter-residue contact maps are shown for the SARS-CoV-2 S complex with H014 - two RBDs in the open state (A), SARS-CoV-2 S complex with H014 - three RBD in the open state (B), SARS-CoV-2 S complex with S309 - two RBDs in the closed form (C), SARS-CoV-2 S complex with S3090 - three RBDs in the closed form (D), SARS-CoV-2 S complex with S2M11 - three RBDs in the closed form (E), SARS-CoV-2 S complex with S2E12 - three RBDs in the closed form (F). The heavy atoms model is employed for computing contacts. The maximum distance for atoms to be considered in the contact is 8 Å

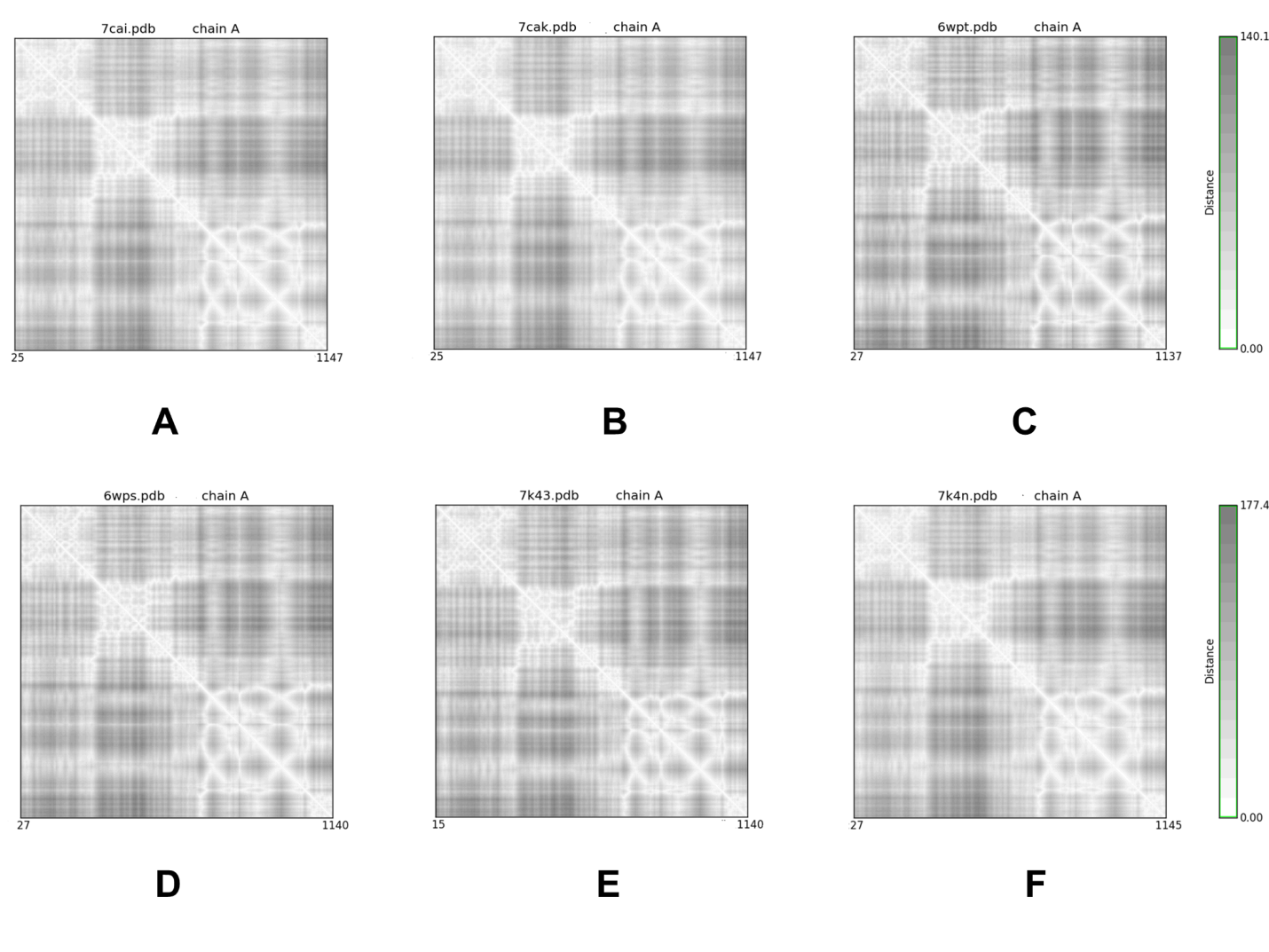
**Figure S5.** The inter-residue distance maps the SARS-CoV-2 S complexes with Abs. (A) The distance maps are shown for the SARS-CoV-2 S complex with H014 - two RBDs in the open state (A), SARS-CoV-2 S complex with H014 - three RBD in the open state (B), SARS-CoV-2 S complex with S309 - two RBDs in the closed form (C), SARS-CoV-2 S complex with S3090 - three RBDs in the closed form (D), SARS-CoV-2 S complex with S2M11 - three RBDs in the closed form (E), SARS-CoV-2 S complex with S2E12 - three RBDs in the closed form (F).

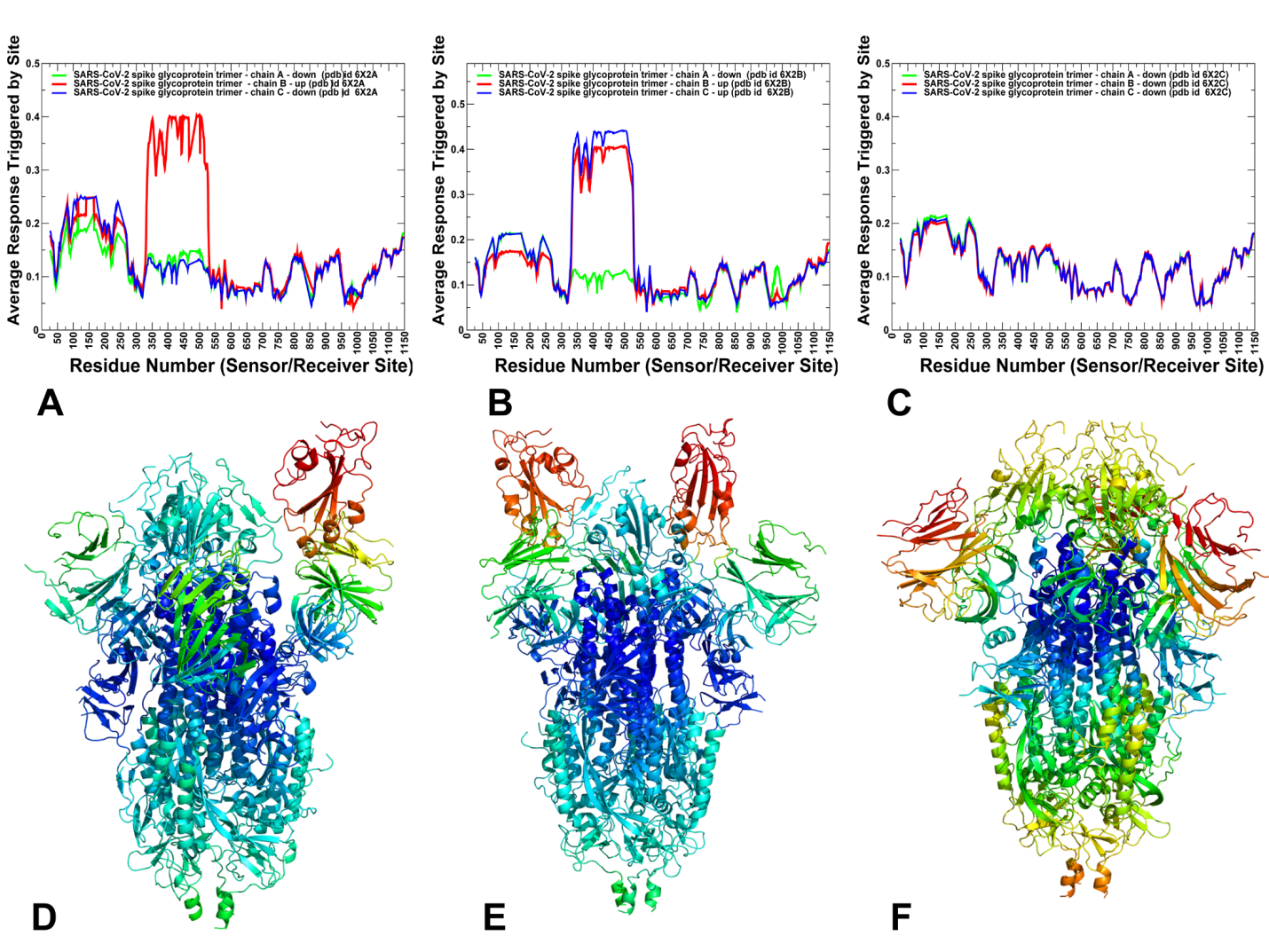
**Figure S6.** The PRS sensor profiles in the closed, partially open and open states of	the SARS-CoV-2 spike trimers. (A) The PRS sensor profile is shown for the ligand-free SARS-CoV-2 S trimer in the partially open (1 RBD up) conformation (pdb id 6X2A). (B) The PRS sensor profile for the ligand-free SARS-CoV-2 S trimer in the	open (2 RBDs up) conformation (pdb id 6X2B). (C)	The PRS sensor profile for the ligand-free SARS-CoV-2 S trimer in the closed (3 RBDs-down) prefusion conformation (pdb id 6X2C). The profiles are shown for protomer in green lines, protomer B in red lines, and protomer C in blue lines. Structural maps of the PRS sensor profiles are shown for the partially open state of the SARS-CoV-2 S prefusion trimer (D), open state (E), and closed state (F). The color gradient from blue to red indicates the increasing sensor propensities. The clusters of residues with the high allosteric potential corresponding to the peaks of the sensor profile.

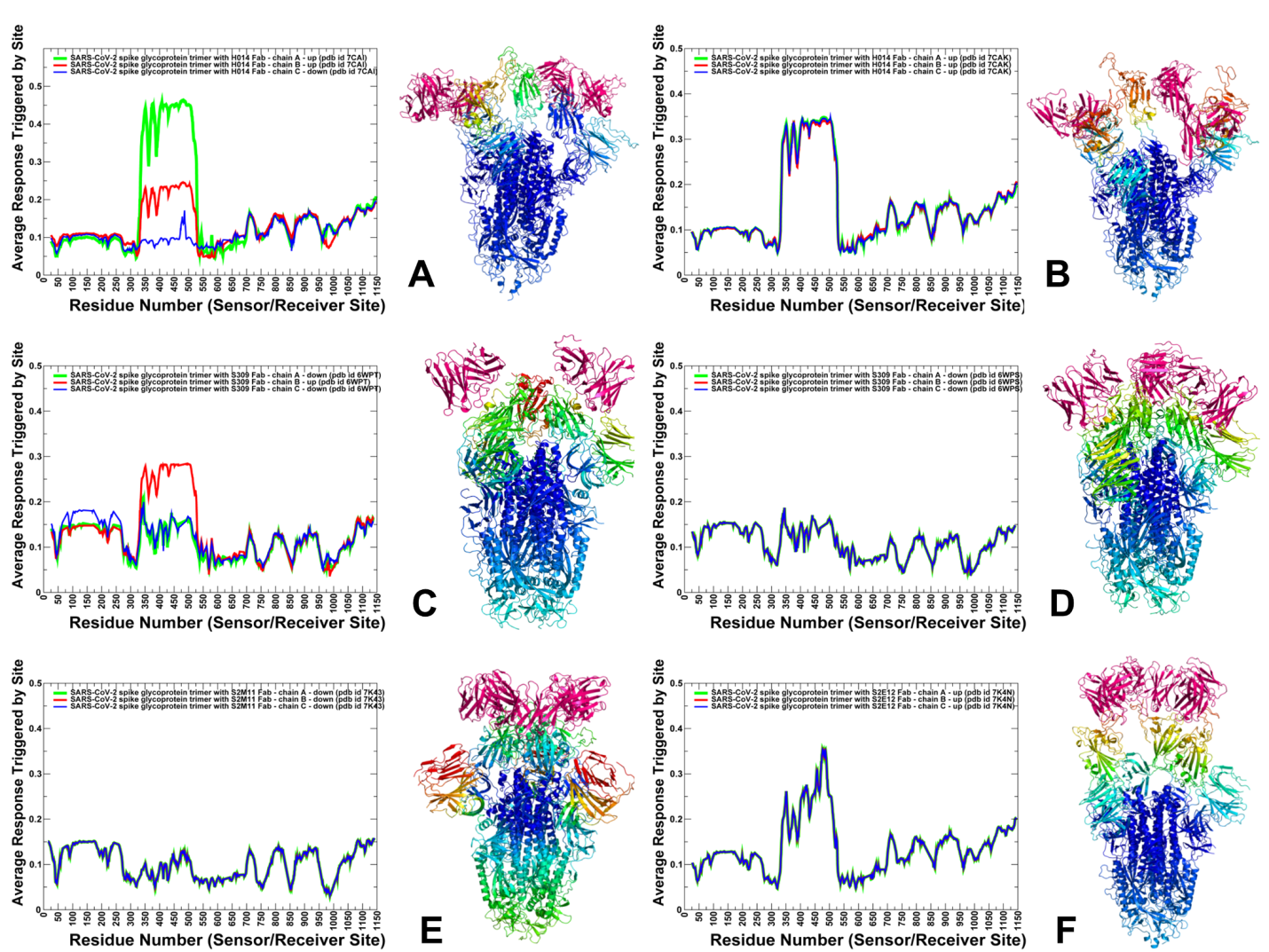
**Figure S7.** The PRS sensor profiles for the SARS-CoV-2 S complexes with Abs. (A) The PRS sensor distributions and structural maps of the sensor profiles are shown for the SARS-CoV-2 S complex with H014 - two RBDs in the open state (A), SARS-CoV-2 S complex with H014 - three RBD in the open state (B), SARS-CoV-2 S complex with S309 - two RBDs in the closed form (C), SARS-CoV-2 S complex with S3090 - three RBDs in the closed form (D), SARS-CoV-2 S complex with S2M11 - three RBDs in the closed form (E), SARS-CoV-2 S complex with S2E12 - three RBDs in the closed form (F). The profiles are shown for protomer in green bars, protomer B in red bars, and protomer C in blue bars. Structural maps of the PRS sensor profiles are shown with the color gradient from blue to red indicating the increasing sensor propensities.

**Table S1.** The list of the intermolecular contacts in the structure of SARS-CoV-2 complex with H014 with 2 protomers in the up state (pdb id 7CAI).

| **S protein residue name** | **S protein residue number** | **S protein chain** | **H014 residue name** | **H014 residue number** | **H014 chain** |
| --- | --- | --- | --- | --- | --- |
| LYS | 378 | A | ASP | 52 | E |
| TYR | 380 | A | TYR | 101 | E |
| ARG | 408 | A | TYR | 101 | E |
| LYS | 378 | A | THR | 58 | E |
| ALA | 411 | A | TYR | 101 | E |
| CYS | 379 | A | ASN | 55 | E |
| PHE | 374 | A | PHE | 92 | D |
| LYS | 378 | A | TYR | 50 | E |
| ARG | 408 | A | PRO | 103 | E |
| GLY | 413 | A | PHE | 54 | E |
| THR | 385 | A | THR | 69 | E |
| GLY | 504 | A | TYR | 104 | E |
| VAL | 382 | A | THR | 58 | E |
| THR | 376 | A | TYR | 105 | E |
| SER | 375 | A | ASN | 91 | D |
| SER | 375 | A | TYR | 95 | D |
| ASP | 427 | A | PHE | 54 | E |
| ASN | 437 | A | SER | 29 | D |
| LYS | 378 | A | TYR | 101 | E |
| LYS | 378 | A | TYR | 33 | E |
| LYS | 378 | A | ILE | 51 | E |
| SER | 373 | A | TRP | 93 | D |
| CYS | 379 | A | THR | 58 | E |
| THR | 376 | A | PRO | 103 | E |
| TYR | 380 | A | TYR | 33 | E |
| LYS | 378 | A | SER | 59 | E |
| SER | 383 | A | GLY | 56 | E |
| VAL | 382 | A | GLY | 57 | E |
| SER | 371 | A | SER | 27 | D |
| ASP | 405 | A | ASP | 102 | E |
| GLN | 414 | A | TYR | 101 | E |
| VAL | 407 | A | ASP | 102 | E |
| TYR | 508 | A | TYR | 104 | E |
| SER | 371 | A | PHE | 92 | D |
| PRO | 384 | A | GLY | 57 | E |
| THR | 385 | A | SER | 59 | E |
| THR | 385 | A | ALA | 68 | E |
| PHE | 377 | A | TYR | 105 | E |
| ARG | 408 | A | ASP | 102 | E |
| PHE | 377 | A | SER | 59 | E |
| TYR | 369 | A | TRP | 93 | D |
| PRO | 384 | A | ASP | 60 | E |
| VAL | 503 | A | SER | 30 | D |
| ASN | 437 | A | ILE | 28 | D |
| PRO | 412 | A | PHE | 54 | E |
| THR | 376 | A | ASN | 91 | D |
| THR | 385 | A | THR | 58 | E |
| TYR | 369 | A | LEU | 62 | E |
| PHE | 374 | A | TRP | 93 | D |
| PRO | 412 | A | ASN | 55 | E |
| THR | 376 | A | TYR | 50 | E |
| PRO | 412 | A | TYR | 101 | E |
| GLN | 506 | A | SER | 30 | D |
| TYR | 380 | A | GLY | 57 | E |
| PHE | 377 | A | TYR | 50 | E |
| LYS | 386 | A | THR | 69 | E |
| SER | 375 | A | TYR | 105 | E |
| GLY | 404 | A | TYR | 104 | E |
| TYR | 380 | A | ASN | 55 | E |
| ASP | 405 | A | TYR | 104 | E |
| LYS | 378 | A | ASN | 55 | E |
| GLY | 404 | A | ASP | 102 | E |
| CYS | 379 | A | GLY | 57 | E |
| TYR | 508 | A | PRO | 103 | E |
| PRO | 384 | A | SER | 59 | E |
| THR | 376 | A | TYR | 33 | E |
| SER | 375 | A | PRO | 103 | E |
| ALA | 372 | A | PHE | 92 | D |
| SER | 383 | A | GLY | 57 | E |
| VAL | 407 | A | PRO | 103 | E |
| ASN | 437 | A | PHE | 92 | D |
| ALA | 372 | A | TRP | 93 | D |
| SER | 383 | A | THR | 69 | E |
| LYS | 378 | A | GLY | 57 | E |
| ALA | 372 | A | SER | 27 | D |
| SER | 373 | A | PHE | 92 | D |
| ASN | 437 | A | ASN | 91 | D |
| THR | 376 | A | ASP | 102 | E |
| TRP | 436 | A | PHE | 92 | D |
| PHE | 374 | A | ASN | 91 | D |
| THR | 385 | A | ASP | 60 | E |
| LEU | 368 | A | TRP | 93 | D |
| SER | 383 | A | THR | 58 | E |
| THR | 385 | A | LYS | 65 | E |
| SER | 375 | A | PHE | 92 | D |
| ALA | 411 | A | ASN | 55 | E |
| PRO | 384 | A | THR | 58 | E |
| CYS | 379 | A | GLY | 56 | E |

**Table S2.** The list of the intermolecular contacts in the structure of SARS-CoV-2 complex with H014 with 3 protomers in the up state (pdb id 7CAK).

| **S protein residue name** | **S protein residue number** | **S protein chain** | **H014 residue name** | **H014 residue number** | **H014 chain** |
| --- | --- | --- | --- | --- | --- |
| GLN | 414 | A | PHE | 54 | E |
| SER | 375 | A | TYR | 105 | E |
| THR | 376 | A | ASN | 91 | D |
| PHE | 377 | A | TYR | 50 | E |
| GLY | 413 | A | TYR | 101 | E |
| GLN | 506 | A | SER | 30 | D |
| ASN | 437 | A | PHE | 92 | D |
| VAL | 503 | A | TYR | 104 | E |
| SER | 375 | A | TYR | 95 | D |
| LYS | 378 | A | SER | 59 | E |
| SER | 375 | A | TRP | 93 | D |
| ARG | 408 | A | TYR | 33 | E |
| PHE | 377 | A | ASP | 60 | E |
| SER | 383 | A | LYS | 65 | E |
| THR | 376 | A | TYR | 50 | E |
| TYR | 369 | A | LEU | 62 | E |
| ASN | 437 | A | ILE | 28 | D |
| ARG | 408 | A | TYR | 101 | E |
| LYS | 378 | A | THR | 58 | E |
| THR | 385 | A | ALA | 67 | E |
| GLY | 413 | A | PHE | 54 | E |
| SER | 375 | A | THR | 90 | D |
| ALA | 411 | A | TYR | 101 | E |
| PRO | 384 | A | SER | 59 | E |
| PRO | 412 | A | TYR | 101 | E |
| ALA | 372 | A | ILE | 2 | D |
| TYR | 380 | A | ASN | 55 | E |
| TRP | 436 | A | PHE | 92 | D |
| GLY | 504 | A | TYR | 104 | E |
| ALA | 372 | A | PHE | 92 | D |
| THR | 376 | A | TYR | 95 | D |
| ALA | 372 | A | SER | 27 | D |
| THR | 376 | A | SER | 59 | E |
| GLN | 414 | A | TYR | 101 | E |
| VAL | 503 | A | SER | 29 | D |
| GLY | 404 | A | PRO | 103 | E |
| ASP | 405 | A | TYR | 104 | E |
| SER | 371 | A | ILE | 2 | D |
| PRO | 384 | A | THR | 58 | E |
| PHE | 374 | A | PHE | 92 | D |
| PRO | 412 | A | PHE | 54 | E |
| THR | 376 | A | TYR | 105 | E |
| SER | 375 | A | PHE | 92 | D |
| VAL | 407 | A | PRO | 103 | E |
| PHE | 374 | A | TRP | 93 | D |
| ALA | 372 | A | TRP | 93 | D |
| PHE | 374 | A | ASN | 91 | D |
| GLN | 414 | A | TYR | 33 | E |
| ASP | 405 | A | ASP | 102 | E |
| SER | 383 | A | ALA | 67 | E |
| ALA | 372 | A | GLN | 26 | D |
| GLN | 414 | A | ASN | 55 | E |
| THR | 385 | A | LEU | 62 | E |
| GLN | 409 | A | ASP | 102 | E |
| ARG | 408 | A | PRO | 103 | E |
| ASP | 427 | A | PHE | 54 | E |
| ALA | 411 | A | ASN | 55 | E |
| CYS | 379 | A | THR | 58 | E |
| PRO | 384 | A | ALA | 68 | E |
| SER | 375 | A | SER | 29 | D |
| LYS | 378 | A | GLY | 56 | E |
| PHE | 374 | A | PRO | 94 | D |
| ALA | 372 | A | GLN | 89 | D |
| TYR | 365 | A | TRP | 93 | D |
| TYR | 380 | A | GLY | 56 | E |
| PHE | 377 | A | SER | 59 | E |
| ASP | 405 | A | PRO | 103 | E |
| VAL | 503 | A | SER | 30 | D |
| VAL | 407 | A | ASP | 102 | E |
| GLN | 506 | A | ASN | 91 | D |
| CYS | 379 | A | GLY | 56 | E |
| GLU | 406 | A | ASP | 102 | E |
| ASN | 437 | A | SER | 29 | D |
| SER | 373 | A | ILE | 2 | D |
| SER | 383 | A | THR | 69 | E |
| PHE | 377 | A | THR | 58 | E |
| TYR | 365 | A | LEU | 62 | E |
| VAL | 433 | A | TYR | 50 | E |
| SER | 375 | A | ASN | 91 | D |
| VAL | 407 | A | TYR | 105 | E |
| THR | 376 | A | PHE | 92 | D |
| GLU | 406 | A | PRO | 103 | E |
| THR | 385 | A | LYS | 65 | E |
| PRO | 384 | A | ASP | 60 | E |
| PHE | 377 | A | TRP | 93 | D |
| TYR | 369 | A | TRP | 93 | D |
| THR | 376 | A | TRP | 93 | D |
| PRO | 384 | A | LYS | 65 | E |
| VAL | 503 | A | ASN | 31 | D |
| SER | 383 | A | ALA | 68 | E |
| LEU | 368 | A | TRP | 93 | D |
| SER | 373 | A | TRP | 93 | D |
| LYS | 378 | A | GLY | 57 | E |
| SER | 373 | A | PHE | 92 | D |
| ARG | 408 | A | ASP | 102 | E |
| PRO | 384 | A | LEU | 62 | E |
| TYR | 380 | A | GLY | 57 | E |
| ASN | 437 | A | ASN | 91 | D |
| TYR | 508 | A | PRO | 103 | E |
| LYS | 378 | A | TYR | 50 | E |
| TYR | 380 | A | THR | 58 | E |
| PRO | 412 | A | ASN | 55 | E |
| TRP | 436 | A | ASN | 91 | D |
| TYR | 508 | A | ASN | 91 | D |

**Table S3.** The list of the intermolecular contacts in the structure of SARS-CoV-2 complex with S309 with 2 protomers in the down state (pdb id 6WPT).

| **S protein residue name** | **S protein residue number** | **S protein chain** | **S309 residue name** | **S309 residue name** | **S309 chain** |
| --- | --- | --- | --- | --- | --- |
| ALA | 344 | A | SER | 109 | D |
| CYS | 361 | A | TRP | 105 | D |
| ARG | 357 | A | PHE | 106 | D |
| LYS | 356 | A | PHE | 106 | D |
| LEU | 335 | A | TYR | 32 | D |
| ASN | 343 | A | ILE | 111 | D |
| THR | 333 | A | THR | 30 | D |
| GLU | 340 | A | GLY | 103 | D |
| ASN | 440 | A | SER | 31 | E |
| GLY | 339 | A | TYR | 100 | D |
| GLU | 340 | A | TYR | 100 | D |
| VAL | 341 | A | PHE | 106 | D |
| ASN | 334 | A | THR | 30 | D |
| ARG | 346 | A | ASP | 93 | E |
| SER | 359 | A | TRP | 105 | D |
| ASN | 343 | A | TYR | 100 | D |
| CYS | 336 | A | TRP | 105 | D |
| ASN | 343 | A | TYR | 50 | E |
| THR | 345 | A | HIS | 92 | E |
| GLU | 340 | A | LEU | 110 | D |
| THR | 345 | A | SER | 33 | E |
| LEU | 335 | A | TRP | 105 | D |
| ALA | 344 | A | GLU | 108 | D |
| ASN | 343 | A | SER | 109 | D |
| GLY | 339 | A | LEU | 110 | D |
| PRO | 337 | A | TRP | 105 | D |
| GLU | 340 | A | ALA | 104 | D |
| ARG | 357 | A | TRP | 105 | D |
| ASN | 360 | A | TYR | 54 | D |
| ARG | 346 | A | SER | 109 | D |
| ALA | 344 | A | ILE | 111 | D |
| ASN | 360 | A | TRP | 105 | D |
| ASN | 334 | A | TYR | 54 | D |
| ILE | 358 | A | PHE | 106 | D |
| GLU | 340 | A | SER | 109 | D |
| LEU | 441 | A | THR | 32 | E |
| LYS | 356 | A | GLU | 108 | D |
| LEU | 441 | A | GLY | 51 | E |
| THR | 333 | A | PHE | 29 | D |
| LEU | 441 | A | SER | 31 | E |
| THR | 345 | A | THR | 32 | E |
| ASN | 343 | A | LEU | 110 | D |
| LEU | 335 | A | SER | 31 | D |
| THR | 345 | A | SER | 109 | D |
| ASN | 334 | A | TRP | 105 | D |
| LEU | 335 | A | THR | 30 | D |
| LEU | 335 | A | PRO | 28 | D |
| ALA | 344 | A | LEU | 110 | D |
| VAL | 341 | A | LEU | 110 | D |
| PRO | 337 | A | PHE | 106 | D |
| ARG | 509 | A | THR | 32 | E |
| THR | 345 | A | LEU | 110 | D |
| SER | 359 | A | TYR | 54 | D |
| GLU | 340 | A | GLU | 108 | D |
| THR | 345 | A | ILE | 111 | D |
| GLU | 340 | A | PHE | 106 | D |
| THR | 333 | A | PRO | 28 | D |

**Table S4.** The list of the intermolecular contacts in the structure of SARS-CoV-2 complex with S309 with 3 protomers in the down state (pdb id 6WPS).

| **S protein residue name** | **S protein residue number** | **S protein chain** | **S309 residue name** | **S309 residue number** | **S309 chain** |
| --- | --- | --- | --- | --- | --- |
| THR | 345 | A | HIS | 92 | L |
| LEU | 335 | A | PRO | 28 | H |
| GLU | 340 | A | ALA | 104 | H |
| ASN | 440 | A | THR | 32 | L |
| THR | 345 | A | ASP | 93 | L |
| ARG | 357 | A | PHE | 106 | H |
| PRO | 337 | A | PHE | 106 | H |
| PRO | 337 | A | TRP | 105 | H |
| GLU | 340 | A | ARG | 102 | H |
| ARG | 346 | A | ASP | 93 | L |
| ASP | 442 | A | SER | 31 | L |
| ILE | 358 | A | PHE | 106 | H |
| CYS | 336 | A | SER | 31 | H |
| LEU | 335 | A | TRP | 105 | H |
| LYS | 356 | A | GLU | 108 | H |
| THR | 345 | A | ILE | 111 | H |
| GLU | 340 | A | GLU | 108 | H |
| ASN | 343 | A | TYR | 100 | H |
| CYS | 361 | A | TRP | 105 | H |
| ASN | 343 | A | ILE | 111 | H |
| ASN | 440 | A | SER | 53 | L |
| VAL | 341 | A | PHE | 106 | H |
| ARG | 509 | A | ILE | 111 | H |
| GLU | 340 | A | TYR | 100 | H |
| LEU | 441 | A | THR | 32 | L |
| THR | 345 | A | THR | 32 | L |
| GLU | 340 | A | GLY | 107 | H |
| LEU | 335 | A | SER | 31 | H |
| ASN | 334 | A | TRP | 105 | H |
| ASN | 354 | A | GLU | 108 | H |
| THR | 333 | A | THR | 30 | H |
| ARG | 346 | A | GLU | 108 | H |
| LEU | 335 | A | THR | 30 | H |
| ARG | 509 | A | THR | 32 | L |
| GLU | 340 | A | GLY | 103 | H |
| ALA | 344 | A | GLU | 108 | H |
| ALA | 344 | A | LEU | 110 | H |
| ASP | 442 | A | THR | 32 | L |
| ARG | 346 | A | SER | 109 | H |
| THR | 345 | A | LEU | 110 | H |
| GLU | 340 | A | LEU | 110 | H |
| THR | 345 | A | SER | 30 | L |
| ALA | 344 | A | SER | 109 | H |
| THR | 345 | A | SER | 33 | L |
| THR | 345 | A | SER | 109 | H |
| ASN | 360 | A | TYR | 54 | H |
| GLU | 340 | A | TRP | 105 | H |
| ASN | 440 | A | SER | 31 | L |
| GLU | 340 | A | SER | 31 | H |
| ASN | 334 | A | TYR | 54 | H |
| THR | 345 | A | THR | 101 | H |
| GLY | 339 | A | LEU | 110 | H |
| ASN | 343 | A | TYR | 50 | L |
| ASN | 360 | A | TRP | 105 | H |
| ASN | 343 | A | SER | 109 | H |
| LEU | 441 | A | SER | 31 | L |
| GLU | 340 | A | PHE | 106 | H |
| SER | 359 | A | TYR | 54 | H |
| SER | 359 | A | TRP | 105 | H |
| ASN | 343 | A | LEU | 110 | H |
| ASN | 334 | A | THR | 30 | H |
| LYS | 356 | A | PHE | 106 | H |
| GLY | 339 | A | TYR | 100 | H |
| ALA | 344 | A | ILE | 111 | H |
| LEU | 441 | A | ILE | 111 | H |
| CYS | 336 | A | TRP | 105 | H |
| ARG | 357 | A | TRP | 105 | H |

**Table S5.** The list of the intermolecular contacts in the structure of SARS-CoV-2 complex with S2M11 with 3 protomers in the down state (pdb id 7K43).

| **S protein residue name** | **S protein residue number** | **S protein chain** | **S2M11 residue name** | **S2M11 residue number** | **S2M11 chain** |
| --- | --- | --- | --- | --- | --- |
| GLN | 498 | A | TYR | 27 | H |
| GLN | 498 | A | THR | 28 | H |
| SER | 373 | A | PHE | 105 | F |
| SER | 494 | A | THR | 28 | H |
| GLN | 493 | A | GLY | 31 | H |
| TYR | 449 | A | THR | 77 | H |
| PHE | 486 | A | PHE | 102 | H |
| CYS | 488 | A | TYR | 33 | H |
| CYS | 488 | A | PHE | 102 | H |
| PHE | 486 | A | SER | 95 | L |
| PHE | 486 | A | TYR | 92 | L |
| PHE | 342 | A | TRP | 106 | F |
| SER | 373 | A | TYR | 103 | F |
| TYR | 449 | A | THR | 30 | H |
| THR | 345 | A | SER | 53 | G |
| TYR | 489 | A | TYR | 103 | H |
| GLU | 484 | A | THR | 58 | H |
| GLU | 484 | A | PHE | 102 | H |
| PHE | 490 | A | ASN | 52 | H |
| GLU | 484 | A | SER | 55 | H |
| PHE | 486 | A | SER | 94 | L |
| TRP | 436 | A | TRP | 106 | F |
| PHE | 486 | A | TYR | 33 | L |
| GLY | 339 | A | PHE | 105 | F |
| ASN | 343 | A | PHE | 105 | F |
| SER | 494 | A | THR | 30 | H |
| GLU | 484 | A | ILE | 54 | H |
| GLY | 485 | A | TYR | 33 | H |
| GLY | 446 | A | THR | 77 | H |
| GLN | 493 | A | THR | 28 | H |
| GLN | 498 | A | GLY | 26 | H |
| LEU | 455 | A | TYR | 103 | H |
| GLN | 493 | A | THR | 30 | H |
| GLU | 484 | A | GLY | 57 | H |
| TYR | 449 | A | THR | 74 | H |
| GLY | 447 | A | THR | 77 | H |
| GLU | 484 | A | SER | 56 | H |
| ASN | 440 | A | THR | 57 | G |
| GLY | 485 | A | TRP | 50 | H |
| GLN | 493 | A | TYR | 32 | H |
| TYR | 449 | A | THR | 28 | H |
| SER | 373 | A | TRP | 106 | F |
| TYR | 495 | A | THR | 28 | H |
| PHE | 490 | A | TYR | 103 | H |
| SER | 371 | A | TYR | 103 | F |
| GLY | 485 | A | PHE | 102 | H |
| SER | 371 | A | PHE | 105 | F |
| LEU | 452 | A | ILE | 54 | H |
| TYR | 489 | A | PHE | 102 | H |
| THR | 345 | A | SER | 54 | G |
| GLY | 446 | A | TYR | 27 | H |
| PHE | 490 | A | ILE | 54 | H |
| VAL | 367 | A | PHE | 105 | F |
| ASN | 440 | A | TYR | 50 | G |
| PHE | 342 | A | PHE | 105 | F |
| VAL | 483 | A | THR | 58 | H |
| LYS | 444 | A | ASP | 61 | G |
| LEU | 441 | A | TYR | 50 | G |
| ALA | 372 | A | TYR | 103 | F |
| SER | 373 | A | ASP | 104 | F |
| TYR | 449 | A | SER | 75 | H |
| TYR | 449 | A | TYR | 27 | H |
| GLU | 484 | A | TYR | 33 | H |
| TYR | 449 | A | PHE | 29 | H |
| LEU | 368 | A | PHE | 105 | F |
| ASN | 343 | A | TRP | 106 | F |
| SER | 494 | A | ILE | 54 | H |
| ASN | 487 | A | PHE | 102 | H |
| LEU | 492 | A | ILE | 54 | H |
| GLU | 484 | A | TRP | 50 | H |
| PHE | 490 | A | SER | 55 | H |
| PHE | 374 | A | TRP | 106 | F |
| GLU | 484 | A | ASN | 52 | H |
| PHE | 456 | A | TYR | 103 | H |
| PHE | 486 | A | TYR | 109 | H |
| GLN | 493 | A | TYR | 103 | H |
| GLY | 447 | A | THR | 74 | H |
| PHE | 374 | A | PHE | 105 | F |
| GLY | 496 | A | THR | 28 | H |
| GLN | 493 | A | ILE | 54 | H |

**Table S6.** The list of the intermolecular contacts in the structure of SARS-CoV-2 complex with S2E12 with 3 protomers in the up state (pdb id 7K4N).

| **S protein residue name** | **S protein residue number** | **S protein chain** | **S2E12 residue name** | **S2E12 residue number** | **S2E12 chain** |
| --- | --- | --- | --- | --- | --- |
| ALA | 475 | A | SER | 105 | H |
| ALA | 475 | A | SER | 107 | H |
| SER | 477 | A | CYS | 106 | H |
| PHE | 486 | A | TYR | 92 | L |
| GLY | 476 | A | CYS | 106 | H |
| GLY | 485 | A | TYR | 92 | L |
| PHE | 486 | A | GLY | 94 | L |
| ALA | 475 | A | CYS | 106 | H |
| LEU | 455 | A | GLY | 54 | H |
| GLN | 474 | A | GLY | 104 | H |
| GLU | 484 | A | GLY | 94 | L |
| LEU | 455 | A | SER | 55 | H |
| SER | 477 | A | ASP | 108 | H |
| ARG | 457 | A | GLY | 104 | H |
| ALA | 475 | A | GLY | 104 | H |
| ALA | 475 | A | SER | 31 | H |
| GLY | 476 | A | ASP | 108 | H |
| THR | 478 | A | SER | 32 | L |
| GLY | 485 | A | LEU | 95 | L |
| ASN | 487 | A | SER | 107 | H |
| GLY | 485 | A | GLY | 94 | L |
| ASN | 487 | A | CYS | 106 | H |
| GLY | 476 | A | SER | 107 | H |
| GLN | 493 | A | SER | 55 | H |
| GLU | 484 | A | LEU | 95 | L |
| PHE | 490 | A | SER | 55 | H |

**Table S7.** The participation coefficients for communicating residues in the structure of SARS-CoV-2 complex with H014 with 2 protomers in the up state (pdb id 7CAI).

| **P** | **Residue Number** | **Chain** | **Residue Name** |
| --- | --- | --- | --- |
| 0.6735 | 322 | {'A'} | {'PRO'} |
| 0.5556 | 323 | {'A'} | {'THR'} |
| 0.7500 | 324 | {'A'} | {'GLU'} |
| 0.6735 | 325 | {'A'} | {'SER'} |
| 0.1736 | 326 | {'A'} | {'ILE'} |
| 0.5556 | 533 | {'A'} | {'LEU'} |
| 0.9600 | 534 | {'A'} | {'VAL'} |
| 0.3056 | 535 | {'A'} | {'LYS'} |
| 0.3600 | 536 | {'A'} | {'ASN'} |
| 0.3056 | 537 | {'A'} | {'LYS'} |
| 0.1736 | 538 | {'A'} | {'CYS'} |
| 0.6213 | 539 | {'A'} | {'VAL'} |
| 0.7500 | 540 | {'A'} | {'ASN'} |
| 0.5556 | 541 | {'A'} | {'PHE'} |
| 0.3600 | 547 | {'A'} | {'THR'} |
| 0.3600 | 548 | {'A'} | {'GLY'} |
| 0.5556 | 549 | {'A'} | {'THR'} |
| 0.5100 | 550 | {'A'} | {'GLY'} |
| 0.1597 | 552 | {'A'} | {'LEU'} |
| 0.2653 | 554 | {'A'} | {'GLU'} |
| 0.1900 | 576 | {'A'} | {'VAL'} |
| 0.5556 | 577 | {'A'} | {'ARG'} |
| 0.5100 | 578 | {'A'} | {'ASP'} |
| 0.2653 | 579 | {'A'} | {'PRO'} |
| 0.6400 | 582 | {'A'} | {'LEU'} |
| 0.8400 | 583 | {'A'} | {'GLU'} |
| 0.3056 | 584 | {'A'} | {'ILE'} |
| 0.3600 | 585 | {'A'} | {'LEU'} |

**Table S8.** The participation coefficients for communicating residues in the structure of SARS-CoV-2 complex with H014 with 3 protomers in the up state (pdb id 7CAK).

| **P** | **Residue Number** | **Chain** | **Residue Name** |
| --- | --- | --- | --- |
| 0.8025 | 296 | {'A'} | {'LEU'} |
| 0.8889 | 307 | {'A'} | {'THR'} |
| 0.7500 | 332 | {'A'} | {'ILE'} |
| 0.7500 | 355 | {'A'} | {'ARG'} |
| 0.7500 | 370 | {'A'} | {'ASN'} |
| 0.7500 | 386 | {'A'} | {'LYS'} |
| 0.7934 | 513 | {'A'} | {'LEU'} |
| 0.7500 | 528 | {'A'} | {'LYS'} |
| 0.8889 | 543 | {'A'} | {'PHE'} |
| 0.7500 | 546 | {'A'} | {'LEU'} |
| 0.7500 | 558 | {'A'} | {'LYS'} |
| 0.7500 | 560 | {'A'} | {'LEU'} |
| 0.8400 | 567 | {'A'} | {'ARG'} |
| 0.8163 | 589 | {'A'} | {'PRO'} |
| 0.8163 | 698 | {'A'} | {'SER'} |
| 0.7500 | 529 | {'B'} | {'LYS'} |
| 0.7500 | 40 | {'C'} | {'ASP'} |
| 0.8163 | 41 | {'C'} | {'LYS'} |
| 0.7500 | 43 | {'C'} | {'PHE'} |
| 0.7500 | 317 | {'C'} | {'ASN'} |
| 0.7500 | 318 | {'C'} | {'PHE'} |
| 0.7500 | 469 | {'C'} | {'SER'} |
| 0.8889 | 470 | {'C'} | {'THR'} |
| 0.8594 | 491 | {'C'} | {'PRO'} |
| 0.8594 | 615 | {'C'} | {'VAL'} |
| 0.8889 | 619 | {'C'} | {'GLU'} |
| 0.8163 | 659 | {'C'} | {'SER'} |
| 0.8025 | 733 | {'C'} | {'LYS'} |
| 0.7500 | 735 | {'C'} | {'SER'} |
| 0.8594 | 863 | {'C'} | {'PRO'} |
| 0.8163 | 866 | {'C'} | {'THR'} |
| 0.9184 | 959 | {'C'} | {'LEU'} |
| 0.8163 | 960 | {'C'} | {'ASN'} |
| 0.7500 | 963 | {'C'} | {'VAL'} |
| 0.9100 | 964 | {'C'} | {'LYS'} |

**Table S9.** The participation coefficients for communicating residues in the structure of SARS-CoV-2 complex with S309 with 2 protomers in the down state (pdb id 6WPT).

| **P** | **Residue Number** | | **Chain** | | **Residue Name** | |
| --- | --- | --- | --- | --- | --- | --- |
| 0.7500 | 558 | | {‘A’} | | {‘LYS’} | |
| 0.7500 | 562 | | {‘A’} | | {‘PHE’} | |
| 0.7500 | 646 | | {‘A’} | | {‘ARG’} | |
| 0.8163 | 659 | | {‘A’} | | {‘SER’} | |
| 0.7500 | 668 | | {‘A’} | | {‘ALA’} | |
| 0.8025 | 697 | | {‘A’} | | {‘MET’} | |
| 0.8889 | 973 | | {‘A’} | | {‘ILE’} | |
| 0.7500 | 987 | | {‘A’} | | {‘PRO’} | |
| 0.7500 | 988 | | {‘A’} | | {‘GLU’} | |
| 0.7500 | 991 | | {‘A’} | | {‘VAL’} | |
| 0.7500 | 40 | | {‘B’} | | {‘ASP’} | |
| 0.7500 | 44 | | {‘B’} | | {‘ARG’} | |
| 0.8594 | 318 | | {‘B’} | | {‘PHE’} | |
| 0.8889 | 319 | | {‘B’} | | {‘ARG’} | |
| 0.8521 | 328 | | {‘B’} | | {‘ARG’} | |
| 0.8025 | 329 | | {‘B’} | | {‘PHE’} | |
| 0.9184 | 335 | | {‘B’} | | {‘LEU’} | |
| 0.9100 | 362 | | {‘B’} | | {‘VAL’} | |
| 0.9184 | 389 | | {‘B’} | | {‘ASP’} | |
| 0.8594 | 527 | | {‘B’} | | {‘PRO’} | |
| 0.9600 | 528 | | {‘B’} | | {‘LYS’} | |
| 0.8889 | 530 | | {‘B’} | | {‘SER’} | |
| 0.8400 | 531 | | {‘B’} | | {‘THR’} | |
| 0.8889 | 592 | | {‘B’} | | {‘PHE’} | |
| 0.8400 | 619 | | {‘B’} | | {‘GLU’} | |
| 0.9600 | 748 | | {‘B’} | | {‘GLU’} | |
| 0.8889 | 855 | | {‘B’} | | {‘PHE’} | |
| 0.8594 | 972 | | {‘B’} | | {‘ALA’} | |
| 0.8400 | 973 | | {‘B’} | | {‘ILE’} | |
| 0.8889 | 994 | | {‘B’} | | {‘ASP’} | |
| 0.7500 | 88 | | {‘C’} | | {‘ASP’} | |
| 0.8678 | 90 | | {‘C’} | | {‘VAL’} | |
| 0.7500 | 194 | | {‘C’} | | {‘PHE’} | |
| 0.7500 | 212 | | {‘C’} | | {‘LEU’} | |
| 0.8400 | 226 | | {‘C’} | | {‘LEU’} | |
| 0.8163 | 264 | | {‘C’} | | {‘ALA’} | |
| 0.8163 | 292 | | {‘C’} | | {‘ALA’} | |
| 0.7500 | 297 | | {‘C’} | | {‘SER’} | |
| 0.7500 | 301 | | {‘C’} | | {‘CYS’} | |
| 0.8163 | 304 | | {‘C’} | | {‘LYS’} | |
| 0.7500 | 553 | | {‘C’} | | {‘THR’} | |
| 0.7500 | 569 | | {‘C’} | | {‘ILE’} | |
| 0.8025 | 571 | | {‘C’} | | {‘ASP’} | |
| 0.7500 | 642 | | {‘C’} | | {‘VAL’} | |
| 0.7500 | 643 | | {‘C’} | | {‘PHE’} | |
| 0.7500 | 653 | | {‘C’} | | {‘ALA’} | |
| 0.7500 | 10 | | {‘E’} | | {‘THR’} | |
| 0.8400 | | 534 | | {‘B’} | | {‘VAL’} |
| 0.9600 | | 535 | | {‘B’} | | {‘LYS’} |
| 0.7934 | | 540 | | {‘B’} | | {‘ASN’} |
| 0.7500 | | 555 | | {‘B’} | | {‘SER’} |
| 0.8025 | | 565 | | {‘B’} | | {‘PHE’} |
| 0.7934 | | 576 | | {‘B’} | | {‘VAL’} |
| 0.7500 | | 43 | | {‘C’} | | {‘PHE’} |
| 0.7500 | | 273 | | {‘C’} | | {‘ARG’} |
| 0.8025 | | 284 | | {‘C’} | | {‘THR’} |
| 0.9100 | | 285 | | {‘C’} | | {‘ILE’} |
| 0.8025 | | 286 | | {‘C’} | | {‘THR’} |

**Table S10.** The participation coefficients for communicating residues in the structure of SARS-CoV-2 complex with S309 with 3 protomers in the down state (pdb id 6WPS).

| **P** | **Residue Number** | **Chain** | **Residue Name** |
| --- | --- | --- | --- |
| 0.7500 | 43 | {'A'} | {'PHE'} |
| 0.7500 | 273 | {'A'} | {'ARG'} |
| 0.7500 | 296 | {'A'} | {'LEU'} |
| 0.8889 | 307 | {'A'} | {'THR'} |
| 0.7500 | 319 | {'A'} | {'ARG'} |
| 0.7500 | 320 | {'A'} | {'VAL'} |
| 0.8025 | 322 | {'A'} | {'PRO'} |
| 0.8889 | 323 | {'A'} | {'THR'} |
| 0.8400 | 342 | {'A'} | {'PHE'} |
| 0.8025 | 363 | {'A'} | {'ALA'} |
| 0.7500 | 364 | {'A'} | {'ASP'} |
| 0.7500 | 365 | {'A'} | {'TYR'} |
| 0.8889 | 366 | {'A'} | {'SER'} |
| 0.7500 | 368 | {'A'} | {'LEU'} |
| 0.8163 | 369 | {'A'} | {'TYR'} |
| 0.7500 | 379 | {'A'} | {'CYS'} |
| 0.9184 | 380 | {'A'} | {'TYR'} |
| 0.8889 | 381 | {'A'} | {'GLY'} |
| 0.8400 | 441 | {'A'} | {'LEU'} |
| 0.7500 | 527 | {'A'} | {'PRO'} |
| 0.9100 | 550 | {'A'} | {'GLY'} |
| 0.8678 | 551 | {'A'} | {'VAL'} |
| 0.7500 | 572 | {'A'} | {'THR'} |
| 0.7500 | 977 | {'A'} | {'LEU'} |
| 0.7500 | 982 | {'A'} | {'SER'} |
| 0.7500 | 983 | {'A'} | {'ARG'} |
| 0.7500 | 988 | {'A'} | {'GLU'} |
| 0.7500 | 39 | {'B'} | {'PRO'} |
| 0.7500 | 49 | {'B'} | {'HIS'} |
| 0.8889 | 57 | {'B'} | {'PRO'} |
| 0.9375 | 58 | {'B'} | {'PHE'} |
| 0.7500 | 272 | {'B'} | {'PRO'} |
| 0.7500 | 288 | {'B'} | {'ALA'} |
| 0.8889 | 319 | {'B'} | {'ARG'} |
| 0.8594 | 320 | {'B'} | {'VAL'} |
| 0.8594 | 334 | {'B'} | {'ASN'} |
| 0.8889 | 335 | {'B'} | {'LEU'} |
| 0.8678 | 336 | {'B'} | {'CYS'} |
| 0.7500 | 344 | {'B'} | {'ALA'} |
| 0.8889 | 346 | {'B'} | {'ARG'} |
| 0.8400 | 358 | {'B'} | {'ILE'} |
| 0.8889 | 373 | {'B'} | {'SER'} |
| 0.7500 | 375 | {'B'} | {'SER'} |
| 0.7500 | 530 | {'B'} | {'SER'} |
| 0.8400 | 531 | {'B'} | {'THR'} |
| 0.7500 | 543 | {'B'} | {'PHE'} |
| 0.8889 | 545 | {'B'} | {'GLY'} |
| 0.7500 | 560 | {'B'} | {'LEU'} |
| 0.7500 | 565 | {'B'} | {'PHE'} |
| 0.7500 | 569 | {'B'} | {'ILE'} |
| 0.8594 | 570 | {'B'} | {'ALA'} |
| 0.8025 | 578 | {'B'} | {'ASP'} |
| 0.9375 | 583 | {'B'} | {'GLU'} |
| 0.8400 | 591 | {'B'} | {'SER'} |
| 0.7500 | 720 | {'B'} | {'ILE'} |
| 0.7500 | 723 | {'B'} | {'THR'} |
| 0.7500 | 891 | {'B'} | {'GLY'} |
| 0.8594 | 902 | {'B'} | {'MET'} |
| 0.7500 | 914 | {'B'} | {'ASN'} |
| 0.8163 | 918 | {'B'} | {'GLU'} |
| 0.8025 | 968 | {'B'} | {'SER'} |
| 0.8025 | 970 | {'B'} | {'PHE'} |
| 0.8594 | 971 | {'B'} | {'GLY'} |
| 0.8889 | 1013 | {'B'} | {'ILE'} |
| 0.7500 | 30 | {'E'} | {'ASN'} |
| 0.7500 | 223 | {'E'} | {'LEU'} |
| 0.8163 | 224 | {'E'} | {'GLU'} |
| 0.8400 | 225 | {'E'} | {'PRO'} |
| 0.7500 | 270 | {'E'} | {'LEU'} |
| 0.7500 | 296 | {'E'} | {'LEU'} |
| 0.7500 | 307 | {'E'} | {'THR'} |
| 0.9722 | 413 | {'E'} | {'GLY'} |
| 0.8889 | 603 | {'E'} | {'ASN'} |
| 0.7500 | 736 | {'E'} | {'VAL'} |
| 0.7500 | 751 | {'E'} | {'ASN'} |
| 0.8025 | 752 | {'E'} | {'LEU'} |
| 0.7500 | 753 | {'E'} | {'LEU'} |
| 0.7500 | 953 | {'E'} | {'ASN'} |
| 0.7500 | 954 | {'E'} | {'GLN'} |
| 0.7500 | 955 | {'E'} | {'ASN'} |
| 0.7500 | 978 | {'E'} | {'ASN'} |
| 0.7500 | 982 | {'E'} | {'SER'} |
| 0.7500 | 983 | {'E'} | {'ARG'} |
| 0.7500 | 1006 | {'E'} | {'THR'} |
| 0.7500 | 1009 | {'E'} | {'THR'} |
| 0.7500 | 1090 | {'E'} | {'PRO'} |
| 0.8400 | 1091 | {'E'} | {'ARG'} |
| 0.8025 | 1103 | {'E'} | {'PHE'} |
| 0.8889 | 1114 | {'E'} | {'ILE'} |
| 0.9100 | 1115 | {'E'} | {'ILE'} |
